# Live-cell imaging of early events following pollen perception in self-incompatible *Arabidopsis thaliana*

**DOI:** 10.1101/760579

**Authors:** Frédérique Rozier, Lucie Riglet, Chie Kodera, Vincent Bayle, Eléonore Durand, Jonathan Schnabel, Thierry Gaude, Isabelle Fobis-Loisy

**Author notes:** Corresponding author: Isabelle Fobis-Loisy, Laboratoire Reproduction et Développement des Plantes, Univ Lyon, ENS de Lyon, UCB Lyon 1, CNRS, INRA, F-69342, Lyon, France. (33) 4 72 72 89 85. Sainsbury Laboratory, University of Cambridge, Bateman Street, Cambridge CB2 1LR, UK. Institut Jean-Pierre Bourgin, Institut National de la Recherche Agronomique (INRA), AgroParisTech, CNRS, Université Paris-Saclay, 78000 Versailles, France.

## Abstract

Early events occurring at the surface of the female organ are critical for plant reproduction, especially in species with a dry stigma. Following landing on the stigmatic papilla cells, the pollen hydrates and germinates a tube, which penetrates the cell wall and grows towards the ovules to convey the male gametes to the embryo sac. In self-incompatible (SI) species within the Brassicaceae, these processes are blocked when the stigma encounters an incompatible pollen. Here, based on the generation of SI-Arabidopsis lines and by setting up a live imaging system, we showed that control of pollen hydration has a central role in pollen selectivity. The faster pollen pumps water from the papilla during an initial period of 10 minutes, the faster it germinates. Furthermore, we found that the SI barriers act to block the proper hydration of incompatible pollen and when hydration is promoted by high humidity, an additional control prevents pollen tube penetration into the stigmatic wall. In papilla cells, actin bundles focalize at the contact site with the compatible pollen but not with the incompatible one, raising the possibility that stigmatic cells react to the mechanical pressure applied by the invading growing tube.

**Highlight:** A live imaging system coupled with self-incompatible Arabidopsis lines highlight the role of stigmatic cells in controlling pollen hydration and in reacting to pollen tube intrusion by remodeling actin cytoskeleton.

## Introduction

Flowers of Brassicaceae species have a dry stigma, highly discriminatory with early control of pollen capture following pollination (Dickinson, 1995). The stigma consists of a dome of flask-shaped epidermal cells (papillae). Once pollen grains land on papillar cells, only those recognized as compatible are accepted, whereas undesirable pollen grains (i.e., from unrelated species or self-pollen in self-incompatible species) are rejected (reviewed in Doucet *et al.* 2016). Pollen grains need to hydrate on the stigma so as to permit the emergence of a pollen tube that penetrates the papilla cell wall and then grows within the pistil tissues (reviewed in Chapman and Goring 2010). *In vitro* assays showed that growing through the stigma is required for pollen tubes to efficiently target the ovules (Palanivelu and Preuss, 2006). Likewise, genetic ablation of the stigmatic papillae prevents normal pollen hydration or germination (Tung *et al.*, 2005). Early events of the pollen-stigma interaction are thus crucial steps for successful fertilization. Most of the Brassicaceae species have a self-incompatibility (SI) system, which allows the stigma to reject self-pollen grains and hence to prevent self-fertilization (de Nettancourt, 2001). This self/non-self recognition mechanism, genetically controlled by a single multiallelic locus, the *S*-locus, depends on a receptor-ligand interaction (Ivanov *et al.*, 2010). The ligand, the *S*-locus Cysteine Rich protein (SCR/SP11), is present in the material contained in the cavities of the pollen cell wall (Schopfer *et al.*, 1999; Takayama *et al.*, 2000), flows out onto the papilla surface and interacts with the *S*-locus Receptor Kinase (SRK) (Stein *et al.*, 1991; Iwano *et al.*, 2003), localized to the plasma membrane of stigmatic papillae. This interaction induces SRK phosphorylation and activation of the signaling cascade leading to self-pollen rejection (Cabrillac *et al.*, 2001). Incompatible pollen grains fail to hydrate properly and do not germinate or only germinate a short pollen tube whose growth is arrested on papilla cells (Dickinson, 1995; Hiroi *et al.*, 2013). Although *Arabidopsis thaliana* is self-fertile, transgenic SI *A. thaliana* lines were generated by reintroducing SCR/SP11 and SRK gene pairs from its close SI relative *A. lyrata* (Nasrallah and Nasrallah, 2002; Nasrallah *et al.*, 2004). SI Arabidopsis lines have opened the way for thorough analysis of pollen-stigma recognition mechanisms through the use of the genetic tools and mutant collections available for this model plant species.

Although they have been studied for many years, the early steps leading to pollen acceptance or rejection decision are still not clearly characterized. In this work, based on the generation of new SI Arabidopsis lines, we revisited the cellular aspects of pollen-stigma interaction by setting up a live imaging system to monitor pollen hydration and germination following compatible as well as incompatible pollinations. We quantified the degree of pollen hydration required for pollen activation and highlighted a late SI control restricting penetration of stigmatic cell wall by the pollen tube. Furthermore, we monitored the dynamics of actin network during compatible and incompatible pollination and found that remodeling of actin cables is triggered in stigmatic cells only when pollen-stigma interaction is engaged in compatibility.

## Materials and methods

### Plant growth conditions

*A. thaliana* Col-0, C24, *A. lyrata ssp petraea* haplotype *S14* originating from the Czech Republic (seeds kindly provided by Dr Pierre Saumitou-Laprade, Université de Lille F59655 Villeneuve d’Ascq cedex, France) and all transgenic plants were grown in growth chambers under a long-day cycle of 16-h light/8-h dark at 21°C/19°C with a relative humidity around 60 %. *A. lyrata* plants were propagated by cuttings every 3 months.

### *A. lyrata SCR14* and *SRK14* gene cloning, plasmid construction, plant transformation and gene expression

The *AlSRK14* genomic sequence spanning the coding region from the start to the stop codons (3,620 kb) and a fragment of 4,081 kb containing the *AlSCR14* gene, 1,844 kb of the 5’ upstream region and 817 bp of the 3’ downstream region were amplified from genomic DNA of *A. lyrata* containing the *S14* locus. Subsequent cloning steps and transgenic lines produced in this work are described in Supplementary protocol S1. Sequences of *AlSRK14* and *AlSCR14* transgenes introduced in *A. thaliana* have been submitted to GENBANK (*pSLR1-gAlSRK14*: accession number MH680585, *pgAlSCR14*: accession number MH680584).

*AlSRK14* expression level in stigma of transgenic plants was determined as described in Supplementary protocol S2.

### Pollination assays

Flower buds at the end of developmental stage 12 (Smyth *et al.*, 1990) were emasculated and 18 hours later stigmas that had reached developmental stage 13 or Early 14 (14E) were manually pollinated with mature pollen. Six hours after pollination, stigmas were fixed in acetic acid 10 %, ethanol 50 % (v/v) and stained with aniline Blue (Kho and Baër, 1968) for pollen tube counting. In a second series of pollination assays, stage 12 flower buds were emasculated and 42h (stage 15) or 66 h (stage 16) later, stigmas were pollinated with fresh mature pollen for 6h followed by fixation and aniline blue staining. In a third experiment, stigmas were pollinated at stage 13-14E and collected after six, 24, 48 or 72 hours followed by fixation and aniline blue staining. Pollination assays were carried out on at least three stigmas and repeated at least at two different dates. Pollination was considered as incompatible when less than five pollen tubes were counted in the stigma (Kitashiba *et al.*, 2011).

### Dual pollination: semi *in vivo* assay

Stage 12 flower bud was emasculated and 18 hours later (stage 13-14E), pistil was cut transversally in the middle of the ovary and placed vertically in a half-cut, perforated PCR tube whose base was introduced in a solid agar medium. Pollen was deposited by gently touching the stigma with a mature anther. Incompatible pollen was deposited first, followed (within less than 1 minute) by deposition of compatible pollen, which defined the timing starting point (T0) for monitoring pollen behavior. A cover slip was delicately applied on the surface of the pollinated stigma for confocal imaging. The system was maintained throughout the experiment at 21°C and under 45 % relative humidity. To increase the relative humidity at the vicinity of the pollinated stigma, pieces of solid agar medium were added around the mounted pistil. For cell surface labelling, after 35 minutes in high humidity conditions, pollinated stigma was incubated in FM™ 4-64 Dye (*N*-3-Triethylammoniumpropyl-4-6-4-Diethylamino Phenyl Hexatrienyl Pyridinium Dibromide, Life Technologies T3166, 8.23 μM) for five minutes and subsequently washed in 1/2 Murashige and Skoog basal medium containing 10 % (w/v) of sucrose, before mounting between slide and coverslip in this medium.

### Confocal microscopy and actin fluorescence intensity

Images were acquired with a SP8 laser-scanning confocal microscope (Leica) using a 25x objective (numeric aperture 0.95, water immersion). Detailed acquisition settings are provided in Supplementary protocol S3. Change in fluorescence intensity in papillae, beneath the pollen grain, around the emerging pollen tube or along the growing tube was estimated by eyes.

### Measurement of pollen hydration rate

Because the increase of water content in pollen grain is correlated with change of its shape from ellipsoid to spheroid (Zuberi and Dickinson, 1985; Hiroi *et al.*, 2013; Wang *et al.*, 2016), for each pollen grain, we recorded every two minutes the ratio between the long and the wide axis (L/W) to estimate the pollen hydration rate until germination. Slopes were determined by linear regression of the hydration curve based on pollen ratio evolution during the first 10 minutes period (5 data points), using the curve f = ax + b. The volume of the pollen grain was estimated using the formula for a prolate spheroid: 4/3 * π * L/2 *(W/2)^2^, where L = length of the longest axis and W = width of the pollen grain.

### Seed set

Flower buds at stage 12 were emasculated and pollinated with mature pollen 18 h later. 21 days after pollination, mature siliques were opened to count the number of seeds.

### Statistic analysis

For each experiment (except the initial selection of SRK and SCR transgenic plants), at least two independent biological replicates (performed at different dates) were carried out. After checking that the mean and variability of the samples were homogenous between replicates, data were pooled to apply a statistical test. Graphs and statistics were obtained with R studio or Excel softwares. We tested the sample distribution with a Shapiro-Wilk test and applied a Student-test to compare the mean of two given data sets. To compare multiple samples with a single test, we performed an ANOVA. In this study, p-values < 0.05 were considered statistically significant.

## Results

### Generation of transgenic lines designed for live cell imaging

We generated transgenic plants for *AlSRK14* in Col-0 background as this accession is more suitable for crossing with available fluorescent marker lines. We used the *Brassica oleracea SLR1* (*S-locus-related gene 1*) promoter to express the *Al*SRK14 receptor in stigmatic cells. Among the 12 transgenic lines we generated, we isolated four independent lines possessing a unique insertion and homozygous for *AlSRK14*. Stage 13-14E stigmas from these four lines rejected pollen from the original *A. lyrata S14* plant (less than five pollen tubes/stigma) whereas *A. lyrata* pollen produced numerous pollen tubes on Col-0 stigmas (Supplementary Fig S1A, B, C). We quantified the expression level of the *AlSRK14* transgene in stage 13-14E stigmas of two of the strongest transgenic plants (lines #10 and #18, Supplementary protocol S2) and found that the relative expression was two times lower than that detected in *A. lyrata S14* stigmas but still sufficient to induce SI response (Supplementary Fig S1D). We selected the transgenic line #10 for further analysis and named it Col-0/*AlSRK14.* In spite of several attempts to introduce *AlSCR14* in Col-0, we did not succeed in generating transgenic pollen that could be rejected by stigmas of Col-0/*AlSRK14*. As an alternative, we used the C24 accession, this accession exhibiting a strong SI response when transformed with *A. lyrata SRK*/*SCR* genes (Nasrallah et al., 2004; Liu et al., 2007). We first transformed C24 plants with *AlSRK14* to generate a female tester capable of rejecting *A. lyrata S14* pollen grains (named C24/*AlSRK14* line #14; Supplementary Fig S1C). Then, among the 18 transgenic lines transformed with *AlSCR14* in C24 background, we isolated five independent lines with unique insertion and homozygous for *AlSCR14*. Pollen from each of these lines was rejected on C24/*AlSRK14* line #14 mature stigmas (Supplementary Fig S1E). We selected transgenic line #4 for further analysis, which was named C24/*AlSCR14.*

To facilitate live-cell monitoring of pollen tube growth on the stigma, we introduced Lifeact fused to Venus to label stigma papillae of Col-0/*AlSRK14 (*Col-0/SRK14 + Act:Venus), TURQUOISE protein to monitor compatible pollen of C24 (C24/TURQ) and RFP for incompatible pollen of C24/*AlSCR14* (C24/SCR14 + RFP) (Fig. 1A). We found that the presence of fluorescent proteins did not alter the compatibility (ANOVA test p-value = 0.23), as well as the SI response (T-test p-value = 0.15) assessed by pollen tube counting (Fig. 1B). Despite the strong pollen rejection response following incompatible cross (female Col-0/SRK14 + Act:Venus x male C24/SCR14 + RFP), some seeds developed in siliques (Fig. 1C). A comparable number of seeds were counted in siliques derived from an incompatible cross between non-labeled pollen and stigmas (mean values of 18.35 versus 19.04, T-test p-value = 0.82, Fig. 1C). However, the incompatible crosses exhibited a significantly reduced seed set compared with the compatible ones (T-test p-value = 2.2E-16 between the four compatible crosses cumulated and the two incompatible crosses cumulated, Fig. 1C). Together, our data show that we succeeded in restoring SI system in *A.thaliana* using a new *A. lyrata* haplotype *S14* and that a mature stigma recognizes and rejects incompatible pollen independently of the accession background. Futhermore, expressing a fluorescent protein in addition to the SI determinant does not modify completion of the SI response.

**Fig. 1.**
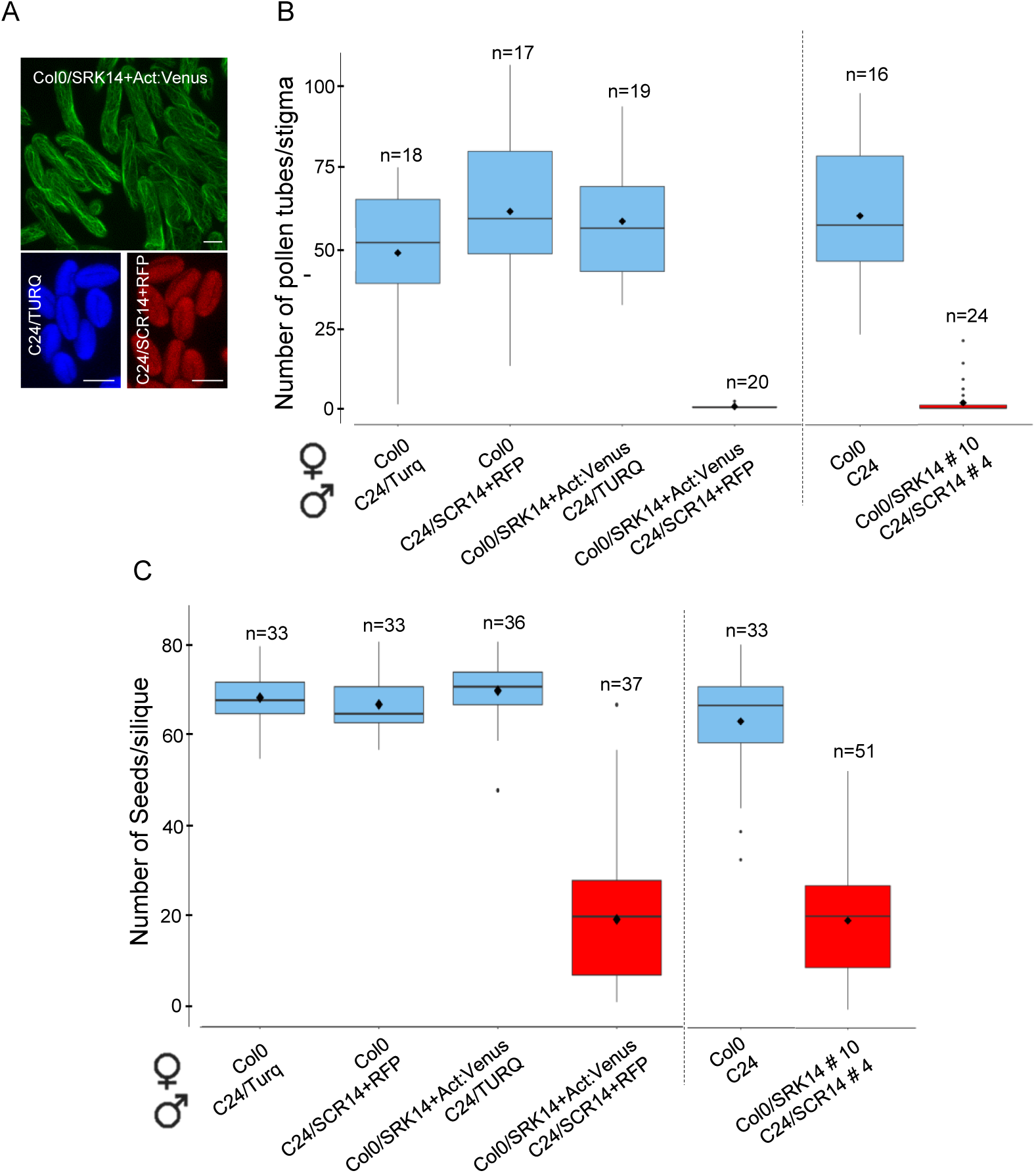
Generation of SI Arabidopsis lines. (A) The Col-0 transgenic line expressing *AlSRK14* was transformed with the fusion protein Act:Venus under the SLR1 promoter to label stigmas (green fluorescence), whereas wild-type C24 or transgenic C24 expressing *AlSCR14* were transformed with the TURQUOISE protein (TURQ, blue fluorescence) or the RFP protein (red fluorescence) under pollen promoters, respectively. Bar = 20 μm. (B) SI phenotype of generated lines. Stage 13-14E stigmas were pollinated with mature pollen and pollen tubes present in stigmas were counted after aniline blue staining. In all crosses, the first name refers to the female partner and the second to the male partner (example: female Col0 x male C24/Turq) (C) Seed set. Stage 13-14E stigmas were pollinated with mature pollen, after 21 days silique were dissected and seeds counted. n = number of pollinated stigmas. Mean (black diamond) of 3 independent experiments. Black dots are extreme values. Experiments separated by a dashed line were performed at different dates. Error bars indicate Standard Error of the Mean (SEM).

### SI response is maintained for about two days and then partially breaks down

As some seeds could develop in siliques following incompatible crosses (Fig. 1C), we investigated how persistent the SI response was during stigma development. To this end, we first carried out pollination experiments using compatible (C24/TURQ) or incompatible (C24/*AlSCR14*) pollen grains deposited on Col-0/SRK14 + Act:Venus stigmas at different pistil developmental stages (Fig. 2A). When incompatible pollen grains were deposited on stage-15 stigmas, almost no pollen tube bypassed the stigmatic barrier (mean of 1.85 tube/stigma, graph Fig. 2A), whereas some pollen tubes succeeded in penetrating stage-16 stigmas (mean of 13.2 tubes/stigma, graph Fig. 2A). By contrast, when compatible pollinations were performed on stigmas at stages 15 or 16, pollen tubes abundantly grew in the stigma. In a second set of experiments, we examined how long incompatible pollen grains deposited on stage 13-14E stigmas could stay inhibited after 6, 24, 48 or 72 hours of contact with papilla cells (Fig. 2B). We observed a breakdown of the SI barrier 48h and 72h after pollen deposition, when stigmas reached stage 16 and stage 17, respectively. However, the number of pollen tubes overpassing SI barrier (mean of 9.9 tubes/stigma and 7.3 tubes/stigma, respectively, graph Fig. 2B) was much less than the above 50 tubes/stigma counted in compatible situations (Graphs Fig. 2A, B). Together, these experiments show that a partial breakdown of the SI response occurs late during pistil ageing and that a strong SI phenotype is maintained during a large time-window of floral development, from stage 13 (anthesis) to stage 15 (stigma extended above long anthers), that is to say slightly less than two days (Smyth *et al.*, 1990). Thus the SI pollination partners we generated are suitable for developing a live-cell imaging to monitor pollen behavior at the surface of mature stigmas (stage 13-14E) without risking weakening of the SI barrier.

**Fig. 2.**
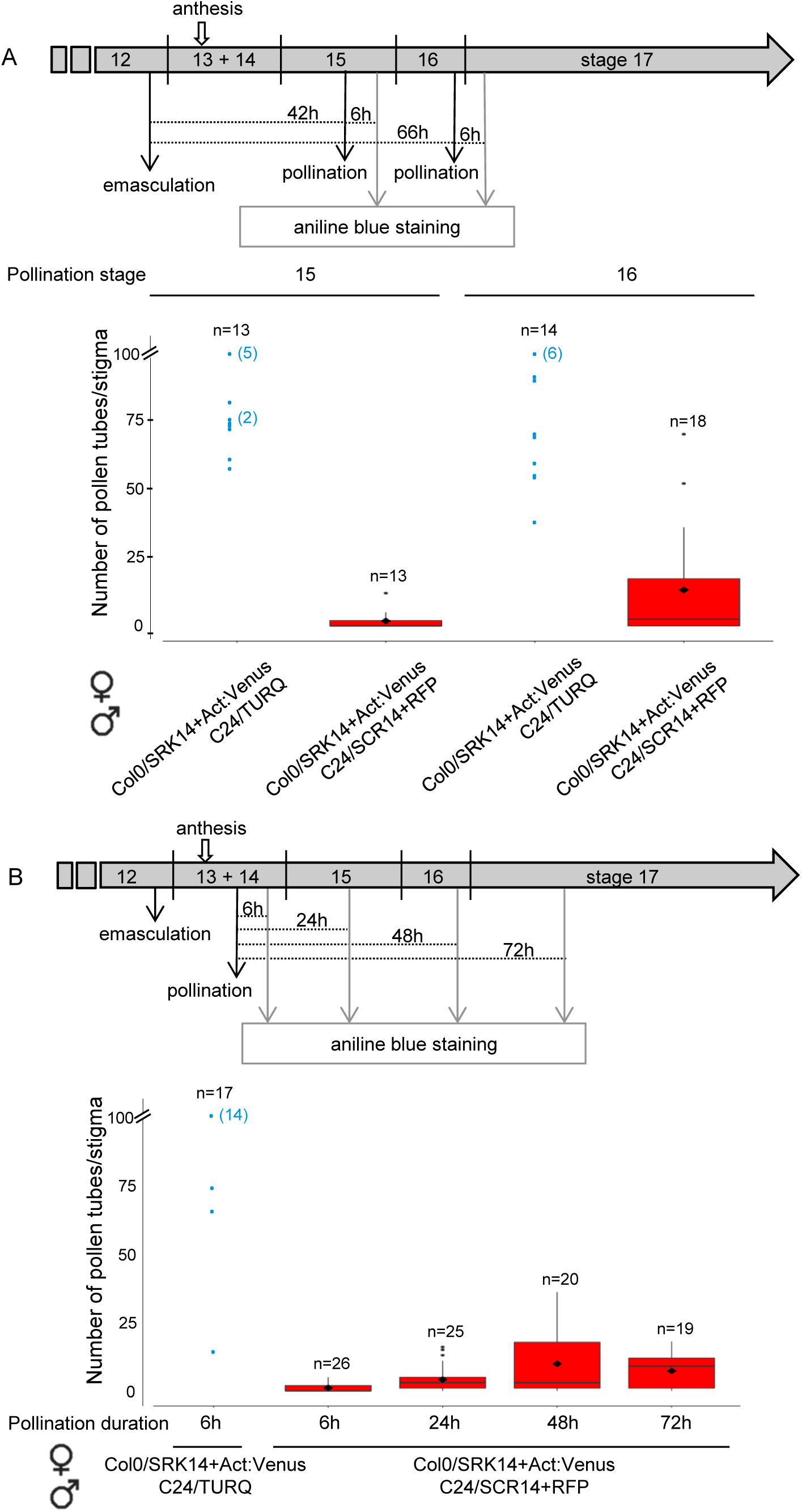
Dynamics of the SI response in Col-0 background. (A) SI phenotype following pollination of stigmas at two developmental stages. Flower buds at the end of developmental stage 12 were emasculated and pollinated with mature pollen 42h (stigma stage 15) or 66h (stigma stage 16) after emasculation (upper panel) and pollen tubes in stigmas were counted 6h after pollination. Pollen tube count after aniline blue staining of stigmas (bottom graph). (B) Maintenance of SI reaction over time. Flower buds at the end of stage 12 were emasculated, pollinated with mature pollen 18h later (stage13-14) and harvested 6, 24, 48 or 72 hours after pollination (upper panel). Pollen tube count after aniline blue staining of stigmas (bottom graph). Beyond 100 pollen tubes per stigma, exact number was not determined and given a value of 100 (no mean, no error bar for compatible crosses). In all crosses, the first name refers to the female partner and the second to the male partner. n = number of pollinated stigmas. Blue dots correspond to individual pollinated stigma; when several stigmas have the same value, the stigma number is indicated in bracket. Mean (black diamond) of two independent experiments. Black dots are extreme values. Error bars indicate SEM.

### Design of a semi *in vivo* assay that allows live cell imaging of early pollination events

To study the very early cellular changes that occur following pollen-stigma interaction, we designed a device that maintains stigma alive for at least one hour and allows cell-live imaging under confocal microscopy (Fig 3A). In this system, compatible and incompatible pollen can be separately deposited on the same stigma and their behavior tracked in the same experimental conditions (Fig 3B, Supplementary video S1). To test whether this setup permitted the maintenance of compatibility/incompatibility responses, we performed five independent experiments where a Col0/SRK14+Act:Venus stigma was dual-pollinated with C24/SCR14+RFP (incompatible) and C24/TURQ (compatible) pollen grains. We found that the percentage of germination for compatible pollen varied from 58 % to 81 %, with a mean value of 74 % (n=106, Fig. 3C). Emission of a pollen tube occurred on average 20 minutes after pollen deposition (Fig. 3C). On the contrary, germination of incompatible pollen was a rare event, occurring for only 1.3 % of the tracked grains (n=75, Fig. 3C). Thus, our semi *in vivo* system allows live cell imaging of pollinated stigmas while preserving the highly regulated SI process.

**Fig. 3.**
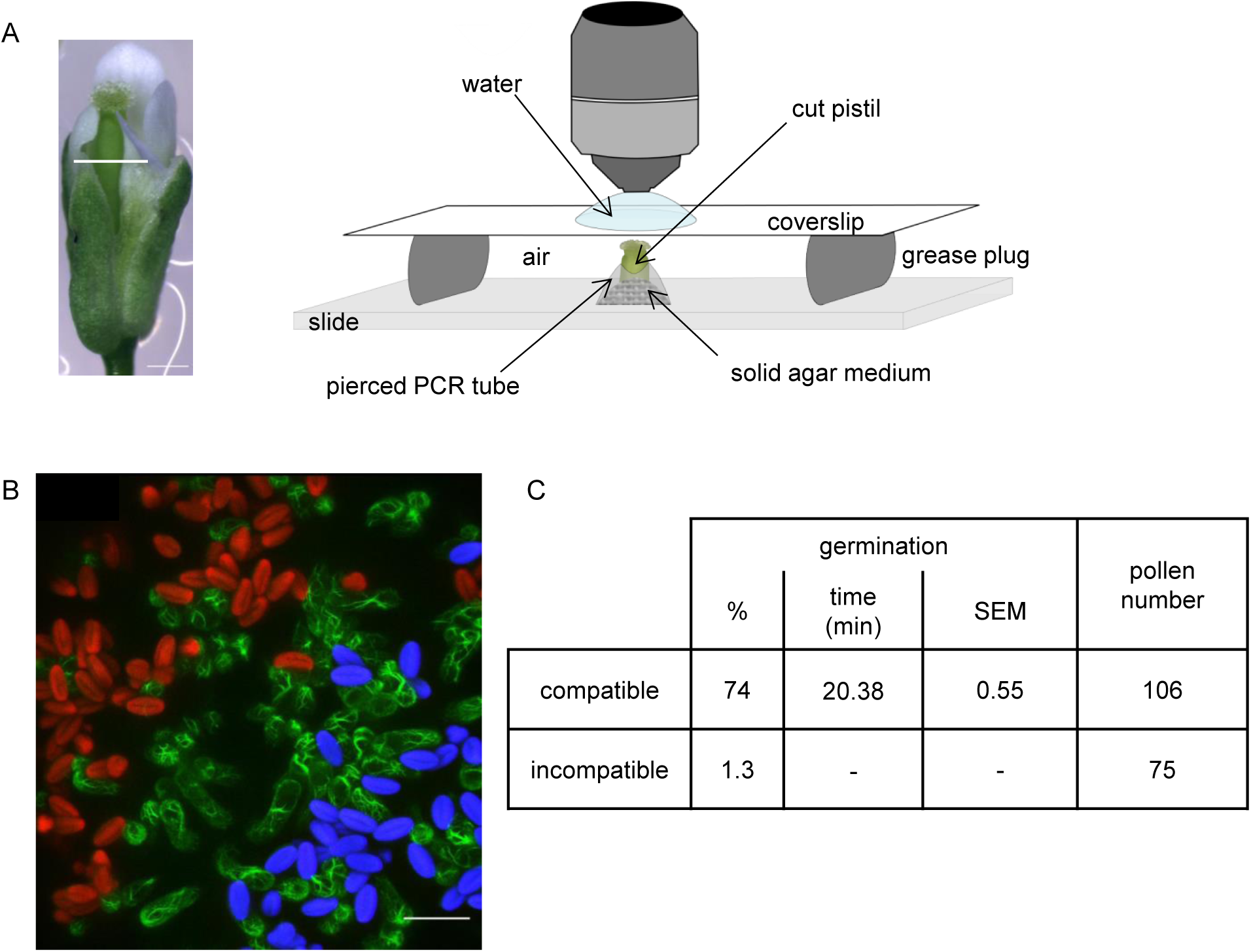
Set up of the semi *in vivo* assay. (A) Upper part of the pistil was cut (white line) and stuck in a solid agar medium. To control humidity level, a small chamber was made with the bottom quarter of a PCR tube whose the base was pierced with a needle and placed upside down on the medium. Stigma was pollinated and a coverslip was gently applied for live imaging. Bar = 500 μm (B) Dual pollination on a single stigma. Stigma (Col-0/SRK14+Act:Venus, green fluorescence) was pollinated first with incompatible pollen (C24/SCR14+RFP, red fluorescence) and immediately after with compatible pollen (C24/TURQ, blue fluorescence), and pollen behavior monitored under confocal microscopy for 32 minutes. Image is a Z-projection (two minutes after pollen deposition). Bar = 50 μm. (C) Pollen germination features extracted from 5 dual-pollinated stigmas. Only one incompatible grain germinated (germination time = 28 minutes). SEM stands for Standard Error of the Mean

### Actin dynamics in stigmatic cells during pollination

Because remodeling of the actin cytoskeleton in the papilla cells has been reported following pollination in Brassica (Iwano *et al.*, 2007), we decided to monitor actin dynamics over time. Before pollination, we found the actin cytoskeleton network to be composed of bundles mainly oriented along the longitudinal axis of the stigmatic cell (Fig. 4A). Upon compatible pollination, a clear accumulation of actin that focalized at the contact site with the compatible pollen was observed in 92 % of the pollinated papillae (22/24, Fig. 4B). For 42% of the stigmatic cells (10/24), actin fluorescence increased beneath the pollen grain just before germination. For 50% of the cells, actin focalized when a clear emergence of the tube (minimum 1 μm) was visible (8%, 2/24) or when the tube started to grow on the stigma surface (42%, 10/24) (Fig. 4B, C, Supplementary video S2, supplementary Fig S3). At later stages of pollen tube progression, actin focalization was observed all along the tube path in 83% (20/24) of the monitored papilla cells (Fig. 4D, Supplementary video S3). Interestingly, when several pollen germinated on the same papilla, actin focalization was observed below each tube (Fig.4E). By contrast, on the seven dual-pollinated stigmas we tracked over time, we never detected any actin focalization beneath the incompatible pollen grains (Supplementary video S4). All together, our live-cell imaging analysis shows that the actin cytoskeleton polymerizes in the stigmatic cell following deposition of compatible but not incompatible pollen grains. Actin accumulation at the pollen contact occurred around the germination process and goes on throughout the elongation of the pollen tube.

**Fig. 4.**
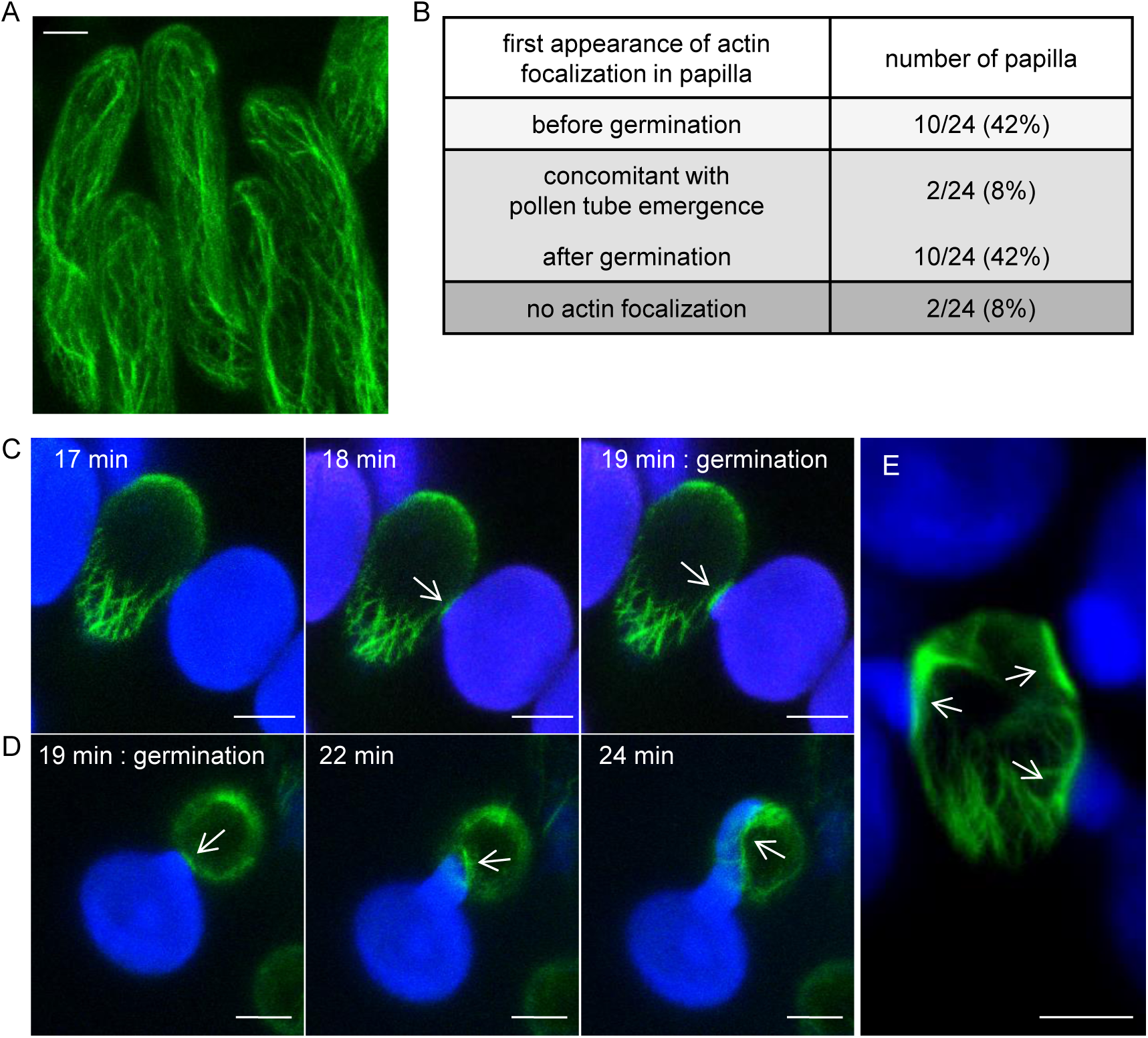
Actin reorganization at the contact site with the compatible pollen grain and pollen tube. (A) Visualization of actin bundles in unpollinated stigmatic cells expressing the Act:Venus marker (Col-0/SRK14+Act:Venus, green fluorescence). Image is a Z-projection. (B) Dynamics of actin focalization following compatible pollination. Act:Venus stigmas were dual-pollinated in the semi *in vivo* system and a Z-stack was taken every minute during 32 minutes. Twenty four Papillae in contact with a compatible pollen grain (blue fluorescence) were selected on 14 stigmas and first appearance of actin fluorescence (focalization) was recovered. Timing of actin focalization was defined regarding the pollen development stage. (C) Actin reorganization in papillae just before pollen germination. (D) Actin reorganization in papillae during germination and pollen tube growth. Indicated time corresponds to time after pollen deposition. Images are Z-projections. (E) Mutiple pollen germination on a single papilla. Image is a Z-projection. Arrows show the focalization of actin fluorescence. Bar = 10 μm.

### Kinetics of compatible pollen hydration is divided into two phases

Next, we compared the kinetics of pollen hydration and germination following dual pollination, using the L/W ratio to estimate the pollen hydration level (Zuberi and Dickinson, 1985). At maturity, the mean L/W ratio of pollen grains was close to two (Fig. 5A, Supplementary table S1). We measured the pollen ratio every two minutes, starting two minutes after pollen deposition (Fig. 5B, Supplementary Fig S2). Among the 69 compatible pollen grains we tracked, 58 germinated in the course of the experiment whereas 11 never did. We found that during the first 10 minutes (preceding germination, Supplementary table S2), the mean pollen ratio dramatically decreased indicating that grains got roundish (solid blue curve Fig. 5B). Interestingly, over the same time period, the pollen volume increased (3455 μm^3^ at 2 min and 4974 μm^3^ at 10 min) strongly supporting that pollen swelling is linked to water uptake and strengthens the use of the L/W ratio as a proxy for pollen hydration. During the following 20 minutes, water uptake almost stopped; indeed, there was no significant difference between the 12 min and 30 min L/W ratios (T-test p-value = 0.71). We found that germination started when the pollen ratio was below 1.4 (solid blue curve Fig. 5B). Among the 58 pollen grains that germinated during the course of the experiment, a huge majority (57/58) reached the 1.4 ratio before germination; only one germinated pollen grain never reached this ratio. For the 11 compatible pollen grains that never germinated (dashed blue curve Fig. 5B), the hydration curve shows no rapid hydration phase during the first 10 minutes and the mean ratio never reached the value of 1.4. Taken together, our results show that in Arabidopsis, water transport toward the pollen grain follows a bi-phasic kinetics: first, an initial phase characterized by a high pollen hydration rate required to reach a certain degree of water content in the pollen (ratio 1.4), followed by a second phase where the hydration rate dramatically decreases and pollen germination occurs.

**Fig. 5.**
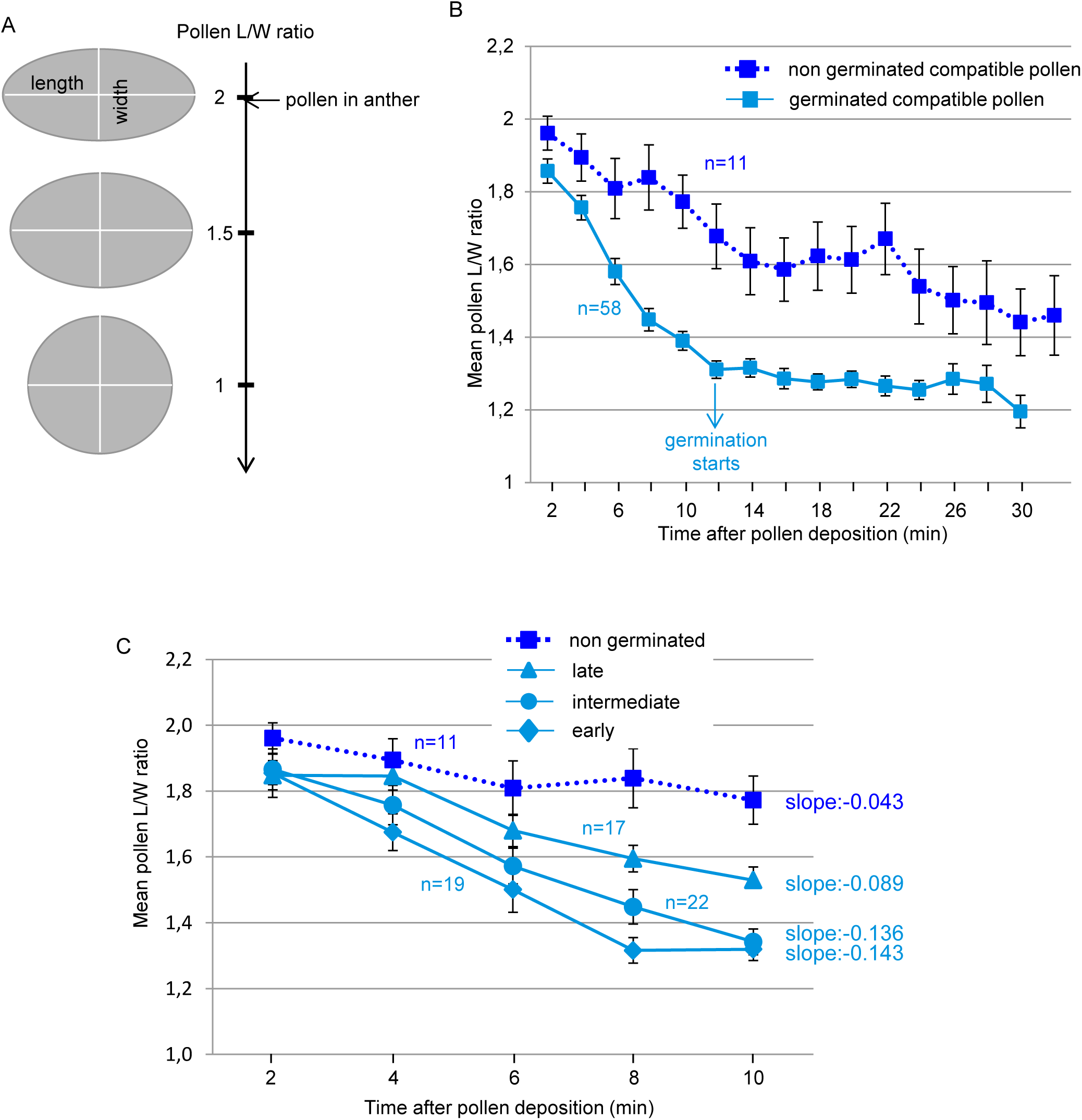
Kinetics of compatible pollen hydration following dual pollination. (A) During hydration, pollen shape changes from an ovoid towards a round shape given by the Length/Width ratio (L/W ratio). (B) Evolution of the L/W ratio during a compatible interaction. Among the 69 tracked pollen grains, 58 germinated during the experiment (solid blue line), whereas 11 failed to germinate (dashed blue line). (C) Kinetics of pollen hydration related to rate of germination. Pollen grains were classified by their germination time in four categories: early, intermediate, late and non germinated. Slopes were calculated by linear regression based on pollen hydration curve during the first 10 minutes. n = number of pollen grains. Mean of five independent experiments. Error bars indicate SEM.

### Variability in pollen germination time depends on the hydration rate during the early interaction phase

In our experimental conditions, pollen germination was not synchronous, starting 12 minutes after pollen contact with the stigma and spreading over the 32 minutes of the experiment (Supplementary table S2). We suspected that different hydration rates of pollen grains might be the cause of this variability. To test this hypothesis, we classified the 69 compatible pollen grains according to their germination capability and categorized them into four classes: those that germinated early (from 12 to 18 minutes, n = 19), intermediate (from 20 to 24 minutes, n = 22), late (from 26 to 32 minutes, n = 17), and those that did not germinate (n = 11), and we compared their hydration rate during the rapid hydration phase from two to 10 minutes (Fig. 5C). At two minutes, no significant differences were found between the L/W ratios of pollen for each classes (1.85/1.87/1.85/1.96, respectively; ANOVA test p-value = 0.13, Fig. 5C). At 10 minutes, pollen that germinated late were significantly less hydrated (higher L/W ratio) than those that germinated early or intermediate (T-test p-value = 2.43E-04 and 1.64E-03, respectively, Fig. 5C), though their hydration degree was significantly higher than non germinated pollen (T-test p-value = 3.62E-02, Fig. 5C). We calculated the slopes of the hydration curves by linear regression, as previously described (Wang *et al.*, 2016), and found that early and intermediate germinating pollen grains had the highest hydration rates compared with late and non germinating pollen grains (Fig 5C). Non germinating pollen exhibited a very slow hydration rate (slope - 0.043) associated with a poor water uptake during the first 10 minutes, deduced from the high pollen L/W ratio at 10 minutes (mean of 1.77). Our data show that most of the tracked compatible pollen germinated within 32 minutes and that the germination time relates to the capacity of the pollen to take water during an initial short period of 10 minutes. The faster the pollen takes water from the stigma, the quicker it germinates.

### Incompatible pollen grains do not fully hydrate and pollen tubes do not penetrate the stigmatic cell wall

Next, we examined the hydration rate of incompatible pollen on the dual-pollinated stigmas. Of the 53 incompatible grains we monitored, none germinated during the time course of the experiment. Futhermore, most of them (n = 43) never exhibited a L/W ratio below the threshold value of 1.4 (solid red line, Fig. 6A) and only hydrated poorly compared with compatible grains (solid blue curve, Fig. 6A). The mean hydration curve for these 43 incompatible grains showed a constant slope during the 32 minutes of the experiment. By contrast, the 10 remaining incompatible grains reached a L/W ratio below 1.4 with a mean hydration curve (dashed red line, Fig. 6A) resembling that of compatible pollen. In addition, their hydration rate within the first 10 minutes was similar to late germinating compatible pollen (slope value of - 0.074 versus - 0.089, respectively, Fig. 6B), and hence hydrated as efficiently as some compatible grains. As high humidity can stimulate pollen hydration and promote incompatible pollen germination (Carter and McNeilly, 1976; Ockendon, 1978; Zuberi and Dickinson, 1985), we checked the effect of high humidity (100 %) on pollen behavior in a modified semi *in vivo* system (Supplementary Fig S3). We found that compatible pollen germination was slightly stimulated compared with standard conditions (88 % germination versus 74 % and mean germination time of 18.55 minutes versus 20.38 minutes, respectively; Supplementary Fig S5). Moreover, high humidity conditions did not dramatically affect actin reorganization as we detected a clear focalization of actin beneath the pollen grain/tube in 75 % of the pollinated papillae (33/44) (Fig. 7A, Supplementary Fig S5, Supplementary video S5). Following incompatible pollination at high humidity, we observed that germination of incompatible pollen was strongly stimulated, varying from 18 % to 58 % depending on the experiment (mean of 35 %, Supplementary Fig S6) compared with the 1.3 % of germination in standard conditions. In addition, among the 46 grains monitored, 46 % reached a L/W ratio of 1.4 within 10 minutes (Supplementary table S3), whereas only 4 % reached this value in standard conditions. However, it took more time for incompatible pollen tubes to emerge compared with compatible tubes (24.87 minutes versus 18.55 minutes, Supplementary Fig S6), and the tube seemed blocked at the papilla surface, the pollen being detached from the papilla (Fig. 7B, Supplementary video S6). The growth rate of incompatible tube was 1.05 μm/minute, whereas a compatible tube extended much more rapidly in both standard and high humidity conditions (2.41 μm/minute and 1.91 μm/minute, respectively, Fig. 7C). Growth kinetics shows that during the first three minutes, elongation rate of both compatible and incompatible tubes was comparable, but then compatible tubes extended rapidly whereas incompatible tubes slowed down (Fig. 7C). These observations suggest that the incompatible tube may not be able to penetrate the papilla cell wall. To test this hypothesis, we stained papilla cells with the amphiphilic styryl dye FM4-64. Contrary to most plant cells where the dye labels the plasma membrane (PM) and the vesicular network (Grebe *et al.*, 2003; Jaillais *et al.*, 2007; Jelínková *et al.*, 2010), we noticed that it remained at the surface of the papilla as it did not colocalize with the actin or the PM canonical marker LTI6b (Supplementary Fig S6). This labelling is probably due to the interaction of the dye with the hydrophobic cuticle layer of the stigmatic cell. While the compatible pollen tube clearly penetrated the FM4-64-labeled layer and grew in close contact with the papilla cytoplasm identified by the actin fluorescence (9 tubes/9, Fig 7D), the incompatible tube was never observed under the FM4-64 layer and stayed outside the papilla (11 tubes/11, Fig 7D). Additionally, at the contact site with the short abnormal incompatible tube, we rarely observed focalization of actin in the papilla, contrary to the compatible situation (Fig. 7B, Supplementary Fig S5 and video S6). Taken together, our results show that incompatible grains generally fail to fully hydrate, and when pollen hydration and germination are promoted by high relative humidity, the SI reaction efficiently blocks tube penetration into the papilla cell wall, the tube remaining outside the papilla surface without triggering actin focalization at the contact site with the stigmatic cell.

**Fig. 6.**
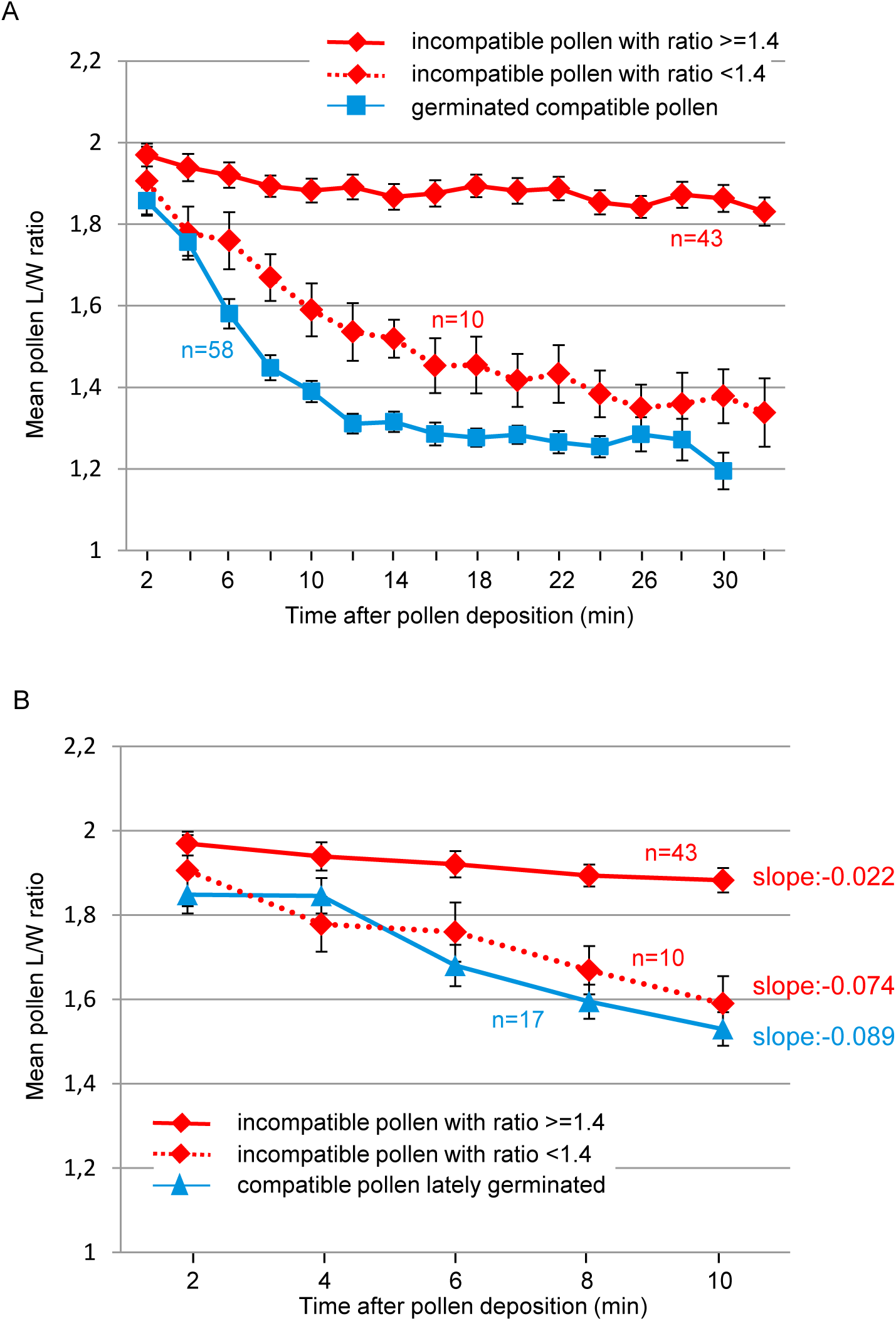
Kinetics of incompatible pollen hydration following dual pollination. (A) Evolution of the L/W ratio during an incompatible interaction. Among the 53 tracked pollen grains, 43 exhibited a L/W ratio that never reached a ratio below 1.4 (solid red line) whereas 10 had a ratio below 1.4 (dashed red line). The solid blue line that indicates hydration dynamics of compatibe pollen was reproduced from Fig. 5B to facilitate the comparison. (B) Hydration rates during the first 10 minutes of incompatible pollen with a L/W ratio >=1.4 (solid red line), incompatible pollen with a L/W ratio <1.4 (dashed red line) and compatible pollen that germinated late (solid blue line, reproduced from Fig. 5C). Slopes were calculated by linear regression. n = number of pollen grains. Mean of 5 independent experiments, error bars indicate SEM.

**Fig. 7.**
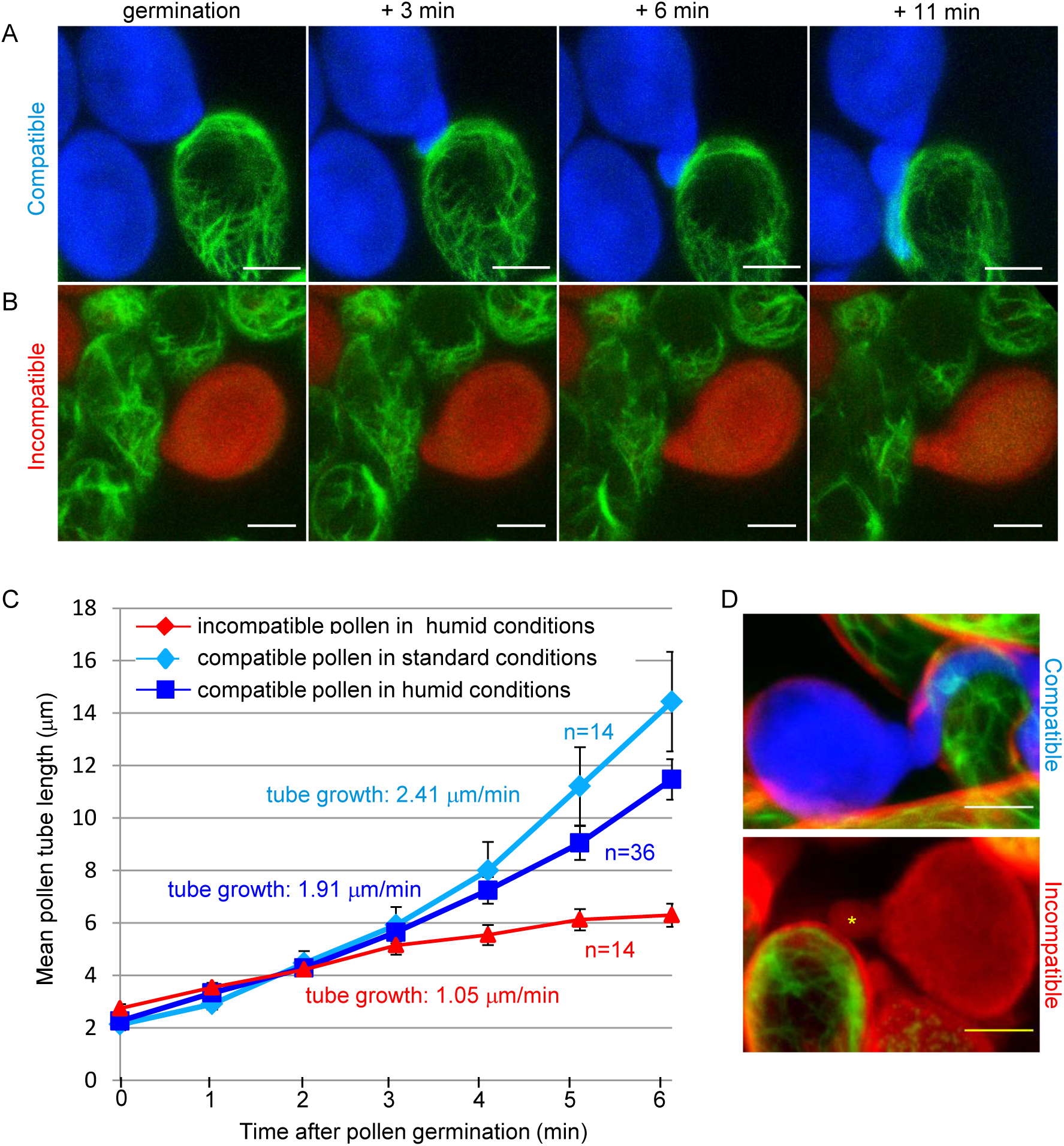
Pollen germination in high humidity conditions. (A) Compatible pollen (blue fluorescence) was deposited on an Act:Venus stigma and incubated in high humidity conditions. (B) Incompatible pollen (red fluorescence) was deposited on an Act:Venus stigma and incubated in high humidity conditions. A Z-stack was taken every minute after pollen deposition. Images are Z-projections. Bar = 10 μm. (C) Growth rate of pollen tubes in standard and high humidity conditions. Pollen tube length was measured at each time point from germination (0 on the horizontal axe) during 6 minutes, for compatible pollen in standard conditions (light blue line), in high humidity conditions (dark blue line) or for incompatible pollen in high humidity conditions (red line). n = number of tracked pollen grains on minimum 6 different stigmas, error bars indicate SEM. (D) 35 minutes after incubation in high humidity conditions, stigmas were stained with FM4-64. A Z-stack was processed with image J to generate a Z-projection. Compatible pollen tube (white star) was detected beneath the FM4-64 labelled layer (red fluorescence surrounding the papilla cell), whereas incompatible tube (yellow star) was outside. Bar = 10 μm.

## Discussion

Here, we set up a live imaging system with a fast-scanning confocal microscope to simultaneously monitor compatible and incompatible Arabidopsis pollen behavior on the same single stigma and provide a thorough analysis of the dynamics of the cellular events following pollen perception.

### *AlSCR14* and *AlSRK14* SI Arabidopsis lines

The two Arabidopsis lines we generated, one that expressed the female determinant (*Al*SRK14) and the second the male determinant (*Al*SCR14) were self-fertile but exhibited a strong SI response when crossed. This SI response was maintained during flower development for about two days, only weakening at late stages of stigma development where few pollen tubes grew in pistil tissues and ultimately led to seed set. Nevertheless, the seed yield remained much below the one reported for selfed Col-0 Arabidopsis (19 seeds per silique versus 62 seeds per silique). Thus, these SRK14 and SCR14 transformants almost fully recapitulate the SI phenotype of the natural SI Brassicaceae, the strength of the response being however less persistent compared with strictly SI lines. Previous work reported a strong SI phenotype, although over a narrow window, in stigmas at stage 13 and early stage 14 in Col-0 lines expressing the cognate *A. lyr*ata *SCRb-SRKb* gene pair (the so-called Col-0/*AlSb* line), whereas older flowers fully accepted self-pollen (Nasrallah, 2002; Nasrallah *et al.*, 2004). By contrast, in another study, the same *SCRb-SRKb* gene pair was shown to be unable to confer SI in Col-0, even in stage-13 flowers, the transgenic plants being fully fertile with a seed set similar to selfed Col-0 plants (Indriolo *et al.*, 2014). In this latter study, a role for the E3-ubiquitin ligase ARC1 in promoting SI was demonstrated as only Col-0 co-expressing the *A. lyrata* or *B. oleracea* ARC1 with the *SCRb-SRKb* pair acquired the capacity to reject self-pollen. Interestingly, the Col-0/*AlSRK14* lines generated in our study express a strong SI response in mature stigmas without the need of co-expressing *AlARC1*, the stigmas remaining partially self-incompatible at later flower developmental stages. Earlier work showed that the variability of SI phenotypes observed between Arabidopsis transformants was mainly due to *SRK* transcript levels (Nasrallah and Nasrallah, 2014). It has been reported that transient SI phenotype and self-fertility of Col-0/*Al*S*b* transformants were caused by modifier loci harbored in the Col-0 genome (Liu *et al.*, 2007; Boggs *et al.*, 2009). One of these loci, *PUB8*, encoding an ARM-repeat and U-box protein, regulates *SRK* transcript levels and is responsible for the pseudo-self compatibility observed in old flowers of Col-0/*AlSb* (Liu *et al.*, 2007). Contrary to the Col-0/*AlSb* transformants mentioned above, in our study, *AlSRK14* expression was driven by the *SLR1* promoter region that does not contain the *SRK* 5’ and 3’ regulatory sequences. As a consequence, it is likely that transcription of *AlSRK14* gene and/or stability of the transcripts escape the regulation of Col-0 SI modifiers, leading to the accumulation of AlSRK14 protein to a level sufficient to maintain a partial pollen rejection response during flower development. The weakening of SI response we observed in late developmental stages was expected as it is known that *SLR1* promoter activity is developmentally regulated, decreasing with flower ageing (Lalonde *et al.*, 1989), and hence leading to lower abundance of *SRK14* transcript levels.

Though we attempted to transform Col-0 with *AlSCR14*, we failed to obtain transformants whose pollen grains were rejected on *AlSRK14*-expressing stigmas. In SI Arabidopsis species, a dominance hierarchy between *SCR* alleles has been reported, which is regulated by small RNAs (sRNAs) (Durand *et al.*, 2014). We may propose that the inability to introduce a functional *AlSCR14* in Col-0 might reside in the presence of vestigial sRNAs targeting the recessive *AlSCR14* allele, while these sRNAs would be absent in the C24 background.

### Pollen behavior

We observed that around 20 minutes after interaction with the stigma papillae, the compatible pollen germinates a tube, which is consistent with the germination time reported in Arabidopsis after *in vivo* pollination (Kandasamy *et al.*, 1994; Iwano, 2004). Once emerged, the pollen tube extends at a rate of 2.41 μm/minute, which is significantly faster than tube elongation *in vitro* (around 1 μm/minute; Iwano *et al.*, 2004; Yang *et al.*, 2017) and consistent with growth rates reported in other *in vivo* systems (Iwano, 2004; Cheung *et al.*, 2010). Futhermore, we found that tube growth rate is not constant, exhibiting a slow growth phase whithin the first three minutes after tube emergence, followed by a second phase of more rapid growth. A bi-phasic growth kinetics was already reported for Arabidopsis pollen tube growing *in vitro*, except that the timing was much longer; the transition from germination to a rapid growth phase requiring 30 minutes (Vogler *et al.*, 2015). While compatible pollen grains promptly achieve germination, incompatible grains deposited on the same stigma, and in the close vicinity of compatible ones, follow a complete different destiny. Indeed, incompatible pollen grains exhibit an extremely low germination rate (1.3 %), which is in agreement with the few pollen tubes we detected after aniline blue staining. Thus, our semi *in vivo* system recapitulates what happens in nature when pollen from various origins lands on the same stigma, accepting compatible whereas rejecting incompatible pollen grains.

We found that almost immediately after landing on the stigma, compatible pollen starts to hydrate and within 10 minutes is almost fully hydrated. By contrast, during the same time period, the L/W ratio of incompatible grains only poorly evolves. Thus, inhibition of the incompatible pollen acts very early, within the first minutes following stigma contact, blocking pollen hydration. As previously described, hydration appears as the first check point controlling pollen rejection in SI species with dry stigma (Zuberi and Dickinson, 1985; Dickinson, 1995; Samuel *et al.*, 2009; Hiroi *et al.*, 2013; Safavian and Goring, 2013; Wang *et al.*, 2016). In this study, the hydration threshold required to trigger germination was found to correspond to a pollen L/W ratio below 1.4. Pollen turgor pressure has been proposed to be the main driving force for germination (Hiroi *et al.*, 2013; Vogler *et al.*, 2015; Wang *et al.*, 2016). Our result suggest that the optimal pollen turgor pressure corresponds to a 1.4 L/W ratio which is reached after 10 minutes of pollen deposition.

Although 19 % (10/53) of the incompatible grains exhibit hydration features similar to the compatible ones, they, however, never germinate. Thus, a second check point acts to ensure that undesirable pollen grains that escape the hydration control are properly inhibited, blocking their germination. When hydration of incompatible pollen is artificially boosted in a saturated humidity environment, a significant proportion of the grains bypasses the second checkpoint and germinates. However, pollen tube growth is slow and does not reach the second phase of rapid elongation observed with compatible tubes. Futhermore, tubes grow outside the papilla cell, unable to penetrate the cell wall. Production of a short incompatible tube, whose growth is arrested in papilla cells, has been described in some *S*-haplotypes of SI Brassica species (Elleman and Dickinson, 1994; Dickinson, 1995). Thus, our data highlight this late control of SI reaction that prevents penetration of the stigmatic cell wall by incompatible tubes. The current SI model in the Brassicaceae proposes that the signaling cascade triggered by SRK activation disrupts the delivery of stigmatic secretory vesicles required for compatible pollen acceptance (reviewed in Jany *et al.*, 2019). The content of these vesicles is currently unknown but regarding the three inhibition levels described in this study and others (Zuberi and Dickinson, 1985; Haasen *et al.*, 2010) we may postulate that stigmatic vesicles transport mainly water to hydrate the pollen grain and trigger germination as well as enzymatic activities to prepare the stigmatic cell wall for pollen tube penetration.

### Actin cytoskeleton

In Brassica, remodeling of the actin cytoskeleton architecture has been described at the site of pollen-stigma interaction (Iwano *et al.*, 2007). However, these results remain controversial, as a previous work suggested that actin filaments do not show any rearrangement (Dearnaley *et al.*, 1999). Using an actin marker line, we show that a clear reorganization of actin occurs at the contact site with compatible pollen. However, while actin bundles appear in Brassica papillae as soon as pollen hydration starts and remain visible during the entire pollen hydration process (Iwano *et al.*, 2007), in Arabidopsis, actin reorganization starts later, from the end of pollen hydration (just before germination) or following pollen tube emergence. One explanation for these discrepancies could be that SI mechanism in Brassica and Arabidopsis does not rely on rigorously similar cellular processes as previously proposed (Kitashiba *et al.*, 2011). Alternatively, these discrepancies could be due to technical reasons: (i) we used stable transformed Arabidopis lines instead of transient expression in *Brassica* papillae, (ii) we performed massive pollination of the stigmatic cells compared with deposition of a single *Brassica* pollen by micromanipulation, and (iii) we used the Lifeact peptide instead of the mouse Talin protein to label F-actin because this short peptide does not affect actin dynamics *in vivo* (Era *et al.*, 2009). Nevertheless, whatever the species, actin focalization appears as a hallmark of the compatible reaction as it is never detected in incompatible pollination. Interestingly, Hardham et al. (2008) showed that touching the surface of the *A. thaliana* cotyledon epidermal cells with a microneedle induced a rapid actin focalization around the contact point, leading the authors to propose that actin reorganization is triggered by detection of mechanical pressure. Likewise, reorganization of actin microfilaments was observed in many plant-pathogen interactions, where actin cables accumulate at the contact site with the pathogen (Takemoto, 2004 and references therein). Altogether, we may hypothetize that reorganization of actin is triggered in stigmatic cells that sense the mechanical pressure produced by the pollen grain/tube at the site of cell wall penetration. In accordance with this, we generally did not detect modification of the actin architecture at the contact site with incompatible pollen tubes that were generated in high humidity conditions and that could not penetrate the stigmatic cell wall. Alternatively, the pollen grain, when engaged in compatibility, could also initiate a signaling cascade leading to the actin polymerization at the contact site with the germinating grain and/or the growing pollen tube. Nevertheless, the focalization of actin is consistent with the well characterized role of actin network in vesicular delivery to the plasma membrane (Szymanski and Staiger, 2018) induced upon compatible pollination.

Taking together, using fast scanning confocal microscopy, our observations underline the suitability of our system to capture very dynamic events occurring within minutes.Notably, we identified a hydration threshold (L/W < 1.4) that pollen grains must reach to acquire the critical water content required for supporting pollen germination. In addition, we monitored the dynamics of actin network during compatible and incompatible responses and found that focalization of actin cables was triggered when pollen-stigma interaction is engaged in compatibility. This actin reorganization might be elicited in response to the mechanical pressure exerted by pollen grain/tube or alternatively by a specific factor delivered by the pollen. Finally, our semi in vivo system provides a particularly well-suited system to test putative components involved in the acceptance or rejection of pollen grains through the analysis of Arabidopsis mutants as well as transgenic lines and will make possible the deciphering of the early events controlling sexual reproduction.

## Supplementary data

**Protocol S1-S3**: supporting material

**Figure S1**: *AlSRK14* and *AlSCR14* Transgenic plant selection

**Video S1**: Dual pollination with compatible and incompatible pollen deposited on the same stigma

**Video S2**: Actin rearrangement at the pollen contact site

**Video S3**: Actin rearrangement along the pollen tube path

**Video S4**: Actin behavior in stigmatic cells in contact with incompatible pollen

**Table S1**: L/W ratio of pollen grains coming out of mature anthers

**Figure S2**: Dynamics of actin focalization following compatible pollination

**Figure S3:** Hydration kinetics of compatible and incompatible pollens in the semi *in vivo* system

**Table S2**: Germination of compatible pollen tracked during experiments described in figure 5

**Figure S4**: Standard and high humidity assays

**Figure S5**: Behavior of compatible pollen in high humidity conditions

**Video S5**: Compatible pollen germination and pollen tube growth in high humidity conditions

**Figure S6**: Behavior of incompatible pollen in high humidity conditions

**Table S3**: L/W ratio of incompatible pollen in high humidity conditions 10 minutes after pollen deposition

**Video S6**: Incompatible pollen germination and pollen tube growth in high humidity conditions

**Figure S7**: FM4-64 labelling of stigmatic cells

## Acknowledgments

We thank Fany Doustaly for *AlSRK14* transgenic line selection, Ghislaine Laroche for *AlSCR14* construction and transgenic line selection, Annie Chaboud for giving us the pAct11 and the pLat52 promoter regions and Claire Lionnet for her support with confocal microscopy. We are grateful to Takashi Ueda for the *LifeActin:Venus* contruct and to Olivier Haman for kindly giving us access to the confocal microscope (ERC-2013-CoG-615739). We thank members of the SiCE team for fruitful discussions. This work was supported by CNRS and Grant ANR-14-CE11-0021.

## References

Boggs NA, Dwyer KG, Nasrallah ME, Nasrallah JB. 2009. In Vivo Detection of Residues Required for Ligand-Selective Activation of the S-Locus Receptor in Arabidopsis. Current Biology 19, 786–791.

Cabrillac D, Cock JM, Dumas C, Gaude T. 2001. The S -locus receptor kinase is inhibited by thioredoxins and activated by pollen coat proteins. Nature 410, 220–223.

Carter AL, McNeilly T. 1976. Increased atmospheric humidity post pollination: A possible aid to the production of inbred line seed from mature flowers in the Brussels sprout (Brassica oleracea var. Gemmifera). Euphytica 25, 531–538.

Chapman LA, Goring DR. 2010. Pollen-pistil interactions regulating successful fertilization in the Brassicaceae. Journal of Experimental Botany 61, 1987–1999.

Cheung AY, Boavida LC, Aggarwal M, Wu H-M, Feijó JA. 2010. The pollen tube journey in the pistil and imaging the in vivo process by two-photon microscopy. Journal of Experimental Botany 61, 1907–1915.

Dearnaley JDW, Clark KM, Heath IB, Lew RR, Goring DR. 1999. Neither compatible nor self-incompatible pollinations of Brassica napus involve reorganization of the papillar cytoskeleton. New phytologist 141, 199–207.

Dickinson H. 1995. Dry stigmas, water and self-incompatibility in Brassica. Sexual Plant Reproduction 8, 1–10.

Doucet J, Lee HK, Goring DR. 2016. Pollen Acceptance or Rejection: A Tale of Two Pathways. Trends in Plant Science 21, 1058–1067.

Durand E, Meheust R, Soucaze M, et al. 2014. Dominance hierarchy arising from the evolution of a complex small RNA regulatory network. Science 346, 1200–1205.

Era A, Tominaga M, Ebine K, Awai C, Saito C, Ishizaki K, Yamato KT, Kohchi T, Nakano A, Ueda T. 2009. Application of Lifeact Reveals F-Actin Dynamics in Arabidopsis thaliana and the Liverwort, Marchantia polymorpha. Plant and Cell Physiology 50, 1041–1048.

Grebe M, Xu J, Möbius W, Ueda T, Nakano A, Geuze HJ, Rook MB, Scheres B. 2003. Arabidopsis sterol endocytosis involves actin-mediated trafficking via ARA6-positive early endosomes. Current Biology 13, 1378–1387.

Haasen KE, Goring DR, others. 2010. The recognition and rejection of self-incompatible pollen in the Brassicaceae. Botanical Studies 51, 1–6.

Hardham AR, Takemoto D, White RG. 2008. Rapid and dynamic subcellular reorganization following mechanical stimulation of Arabidopsis epidermal cells mimics responses to fungal and oomycete attack. BMC Plant Biology 8, 63.

Hiroi K, Sone M, Sakazono S, Osaka M, Masuko-Suzuki H, Matsuda T, Suzuki G, Suwabe K, Watanabe M. 2013. Time-lapse imaging of self- and cross-pollinations in Brassica rapa. Annals of Botany 112, 115–122.

Indriolo E, Safavian D, Goring DR. 2014. The ARC1 E3 Ligase Promotes Two Different Self-Pollen Avoidance Traits in Arabidopsis. The Plant Cell 26, 1525–1543.

Ivanov R, Fobis-Loisy I, Gaude T. 2010. When no means no: guide to Brassicaceae self-incompatibility. Trends in Plant Science 15, 387–394.

Iwano M. 2004. Ca2+ Dynamics in a Pollen Grain and Papilla Cell during Pollination of Arabidopsis. PLANT PHYSIOLOGY 136, 3562–3571.

Iwano M, Shiba H, Funato M, Shimosato H, Takayama S, Isogai A. 2003. Immunohistochemical studies on translocation of pollen S-haplotype determinant in self-incompatibility of Brassica rapa. Plant and cell physiology 44, 428–436.

Iwano M, Shiba H, Matoba K, et al. 2007. Actin Dynamics in Papilla Cells of Brassica rapa during Self- and Cross-Pollination. PLANT PHYSIOLOGY 144, 72–81.

Jaillais Y, Fobis-Loisy I, Miège C, Gaude T. 2007. Evidence for a sorting endosome in Arabidopsis root cells: Plant sorting endosome. The Plant Journal 53, 237–247.

Jany E, Nelles H, Goring DR. 2019. The Molecular and Cellular Regulation of Brassicaceae Self-Incompatibility and Self-Pollen Rejection. International Review of Cell and Molecular Biology. Elsevier, 1–35.

Jelínková A, Malínská K, Simon S, et al. 2010. Probing plant membranes with FM dyes: tracking, dragging or blocking? The Plant Journal 61, 883–892.

Kandasamy MK, Nasrallah JB, Nasrallah ME. 1994. Pollen-pistil interactions and developmental regulation of pollen tube growth in Arabidopsis. Development 120, 3405–3418.

Kho YO. Observing pollen tubes by means of fluorescence., 5.

Kitashiba H, Liu P, Nishio T, Nasrallah JB, Nasrallah ME. 2011. Functional test of Brassica self-incompatibility modifiers in Arabidopsis thaliana. Proceedings of the National Academy of Sciences 108, 18173–18178.

Lalonde BA, Nasrallah ME, Dwyer KG, Chen C-H, Barlow B, Nasrallah JB. 1989. A highly conserved Brassica gene with homology to the S-locus-specific glycoprotein structural gene. The Plant Cell 1, 249–258.

Liu P, Sherman-Broyles S, Nasrallah ME, Nasrallah JB. 2007. A Cryptic Modifier Causing Transient Self-Incompatibility in Arabidopsis thaliana. Current Biology 17, 734–740.

Nasrallah ME. 2002. Generation of Self-Incompatible Arabidopsis thaliana by Transfer of Two S Locus Genes from A. lyrata. Science 297, 247–249.

Nasrallah ME, Liu P, Sherman-Broyles S, Boggs NA, Nasrallah JB. 2004. Natural variation in expression of self-incompatibility in Arabidopsis thaliana: implications for the evolution of selfing. Proceedings of the National Academy of Sciences of the United States of America 101, 16070–16074.

Nasrallah JB, Nasrallah ME. 2014. *S* -locus receptor kinase signalling. Biochemical Society Transactions 42, 313–319.

Nettancourt D de. 2001. Incompatibility and Incongruity in Wild and Cultivated Plants. Berlin Heidelberg: Springer-Verlag.

Ockendon DJ. 1978. Effect of hexane and humidity on self-incompatibility in Brassica oleracea. Theoretical and Applied Genetics 52, 113–117.

Palanivelu R, Preuss D. 2006. Distinct short-range ovule signals attract or repel Arabidopsis thaliana pollen tubes in vitro. BMC Plant Biology 6, 7.

Safavian D, Goring DR. 2013. Secretory Activity Is Rapidly Induced in Stigmatic Papillae by Compatible Pollen, but Inhibited for Self-Incompatible Pollen in the Brassicaceae (D Bassham, Ed.). PLoS ONE 8, e84286.

Samuel MA, Chong YT, Haasen KE, Aldea-Brydges MG, Stone SL, Goring DR. 2009. Cellular Pathways Regulating Responses to Compatible and Self-Incompatible Pollen in Brassica and Arabidopsis Stigmas Intersect at Exo70A1, a Putative Component of the Exocyst Complex. THE PLANT CELL ONLINE 21, 2655–2671.

Schopfer CR. 1999. The Male Determinant of Self-Incompatibility in Brassica. Science 286, 1697–1700.

Smyth DR, Bowman JL, Meyerowitz EM. 1990. Early flower development in Arabidopsis. The Plant Cell 2, 755–767.

Stein JC, Howlett B, Boyes DC, Nasrallah ME, Nasrallah JB. 1991. Molecular cloning of a putative receptor protein kinase gene encoded at the self-incompatibility locus of Brassica oleracea. Proceedings of the National Academy of Sciences 88, 8816–8820.

Szymanski D, Staiger CJ. 2018. The Actin Cytoskeleton: Functional Arrays for Cytoplasmic Organization and Cell Shape Control. Plant Physiology 176, 106–118.

Takayama S, Shiba H, Iwano M, Shimosato H, Che F-S, Kai N, Watanabe M, Suzuki G, Hinata K, Isogai A. 2000. The pollen determinant of self-incompatibility in Brassica campestris. Proceedings of the National Academy of Sciences 97, 1920–1925.

Takemoto D. 2004. The Cytoskeleton as a Regulator and Target of Biotic Interactions in Plants. PLANT PHYSIOLOGY 136, 3864–3876.

Tung C-W. 2005. Genome-Wide Identification of Genes Expressed in Arabidopsis Pistils Specifically along the Path of Pollen Tube Growth. PLANT PHYSIOLOGY 138, 977–989.

Vogler F, Konrad SSA, Sprunck S. 2015. Knockin’ on pollen’s door: live cell imaging of early polarization events in germinating Arabidopsis pollen. Frontiers in Plant Science 6.

Wang L, Clarke LA, Eason RJ, Parker CC, Qi B, Scott RJ, Doughty J. 2016. PCP-B class pollen coat proteins are key regulators of the hydration checkpoint in *Arabidopsis thaliana* pollen-stigma interactions. New Phytologist.

Yang X, Zhang Q, Zhao K, Luo Q, Bao S, Liu H, Men S. 2017. The Arabidopsis GPR1 Gene Negatively Affects Pollen Germination, Pollen Tube Growth, and Gametophyte Senescence. International Journal of Molecular Sciences 18.

Zuberi MI, Dickinson HG. 1985. Pollen-stigma interaction in Brassica. III. Hydration of the pollen grains. Journal of Cell Science 76, 321–336.

