## Supplementary data for "Live-cell imaging of early events following pollen perception in self-incompatible *Arabidopsis thaliana*"

### Slide 1
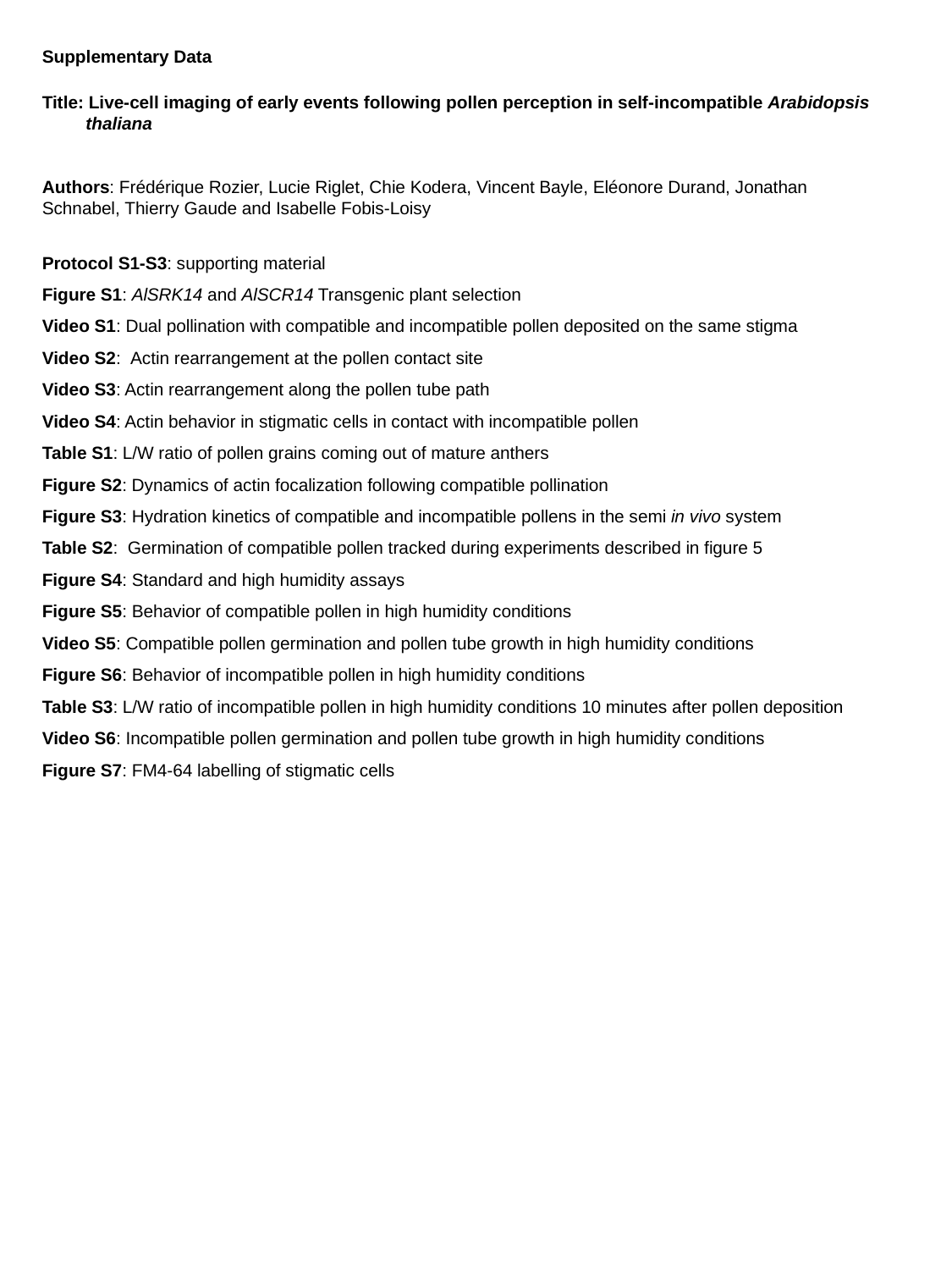

Supplementary Data
Title: Live-cell imaging of early events following pollen perception in self-incompatible Arabidopsis
 thaliana
Authors: Frédérique Rozier, Lucie Riglet, Chie Kodera, Vincent Bayle, Eléonore Durand, Jonathan Schnabel, Thierry Gaude and Isabelle Fobis-Loisy
Protocol S1-S3: supporting material
Figure S1: AlSRK14 and AlSCR14 Transgenic plant selection
Video S1: Dual pollination with compatible and incompatible pollen deposited on the same stigma
Video S2: Actin rearrangement at the pollen contact site
Video S3: Actin rearrangement along the pollen tube path
Video S4: Actin behavior in stigmatic cells in contact with incompatible pollen
Table S1: L/W ratio of pollen grains coming out of mature anthers
Figure S2: Dynamics of actin focalization following compatible pollination
Figure S3: Hydration kinetics of compatible and incompatible pollens in the semi in vivo system
Table S2: Germination of compatible pollen tracked during experiments described in figure 5
Figure S4: Standard and high humidity assays
Figure S5: Behavior of compatible pollen in high humidity conditions
Video S5: Compatible pollen germination and pollen tube growth in high humidity conditions
Figure S6: Behavior of incompatible pollen in high humidity conditions
Table S3: L/W ratio of incompatible pollen in high humidity conditions 10 minutes after pollen deposition
Video S6: Incompatible pollen germination and pollen tube growth in high humidity conditions
Figure S7: FM4-64 labelling of stigmatic cells

### Slide 2
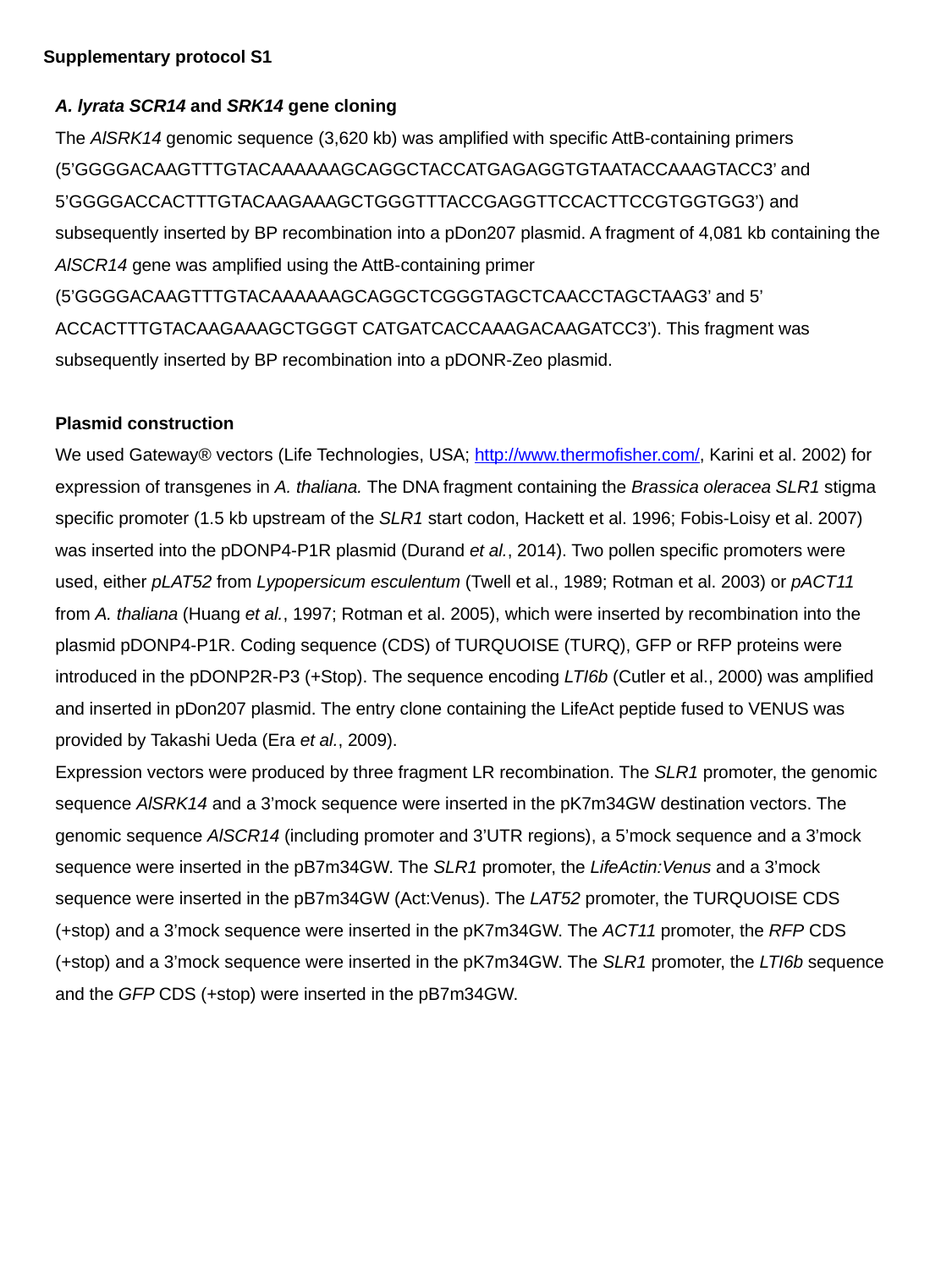

Supplementary protocol S1
A. lyrata SCR14 and SRK14 gene cloning
The AlSRK14 genomic sequence (3,620 kb) was amplified with specific AttB-containing primers (5’GGGGACAAGTTTGTACAAAAAAGCAGGCTACCATGAGAGGTGTAATACCAAAGTACC3’ and 5’GGGGACCACTTTGTACAAGAAAGCTGGGTTTACCGAGGTTCCACTTCCGTGGTGG3’) and subsequently inserted by BP recombination into a pDon207 plasmid. A fragment of 4,081 kb containing the AlSCR14 gene was amplified using the AttB-containing primer (5’GGGGACAAGTTTGTACAAAAAAGCAGGCTCGGGTAGCTCAACCTAGCTAAG3’ and 5’ ACCACTTTGTACAAGAAAGCTGGGT CATGATCACCAAAGACAAGATCC3’). This fragment was subsequently inserted by BP recombination into a pDONR-Zeo plasmid.
Plasmid construction
We used Gateway® vectors (Life Technologies, USA; http://www.thermofisher.com/, Karini et al. 2002) for expression of transgenes in A. thaliana. The DNA fragment containing the Brassica oleracea SLR1 stigma specific promoter (1.5 kb upstream of the SLR1 start codon, Hackett et al. 1996; Fobis-Loisy et al. 2007) was inserted into the pDONP4-P1R plasmid (Durand et al., 2014). Two pollen specific promoters were used, either pLAT52 from Lypopersicum esculentum (Twell et al., 1989; Rotman et al. 2003) or pACT11 from A. thaliana (Huang et al., 1997; Rotman et al. 2005), which were inserted by recombination into the plasmid pDONP4-P1R. Coding sequence (CDS) of TURQUOISE (TURQ), GFP or RFP proteins were introduced in the pDONP2R-P3 (+Stop). The sequence encoding LTI6b (Cutler et al., 2000) was amplified and inserted in pDon207 plasmid. The entry clone containing the LifeAct peptide fused to VENUS was provided by Takashi Ueda (Era et al., 2009).
Expression vectors were produced by three fragment LR recombination. The SLR1 promoter, the genomic sequence AlSRK14 and a 3’mock sequence were inserted in the pK7m34GW destination vectors. The genomic sequence AlSCR14 (including promoter and 3’UTR regions), a 5’mock sequence and a 3’mock sequence were inserted in the pB7m34GW. The SLR1 promoter, the LifeActin:Venus and a 3’mock sequence were inserted in the pB7m34GW (Act:Venus). The LAT52 promoter, the TURQUOISE CDS (+stop) and a 3’mock sequence were inserted in the pK7m34GW. The ACT11 promoter, the RFP CDS (+stop) and a 3’mock sequence were inserted in the pK7m34GW. The SLR1 promoter, the LTI6b sequence and the GFP CDS (+stop) were inserted in the pB7m34GW.

### Slide 3
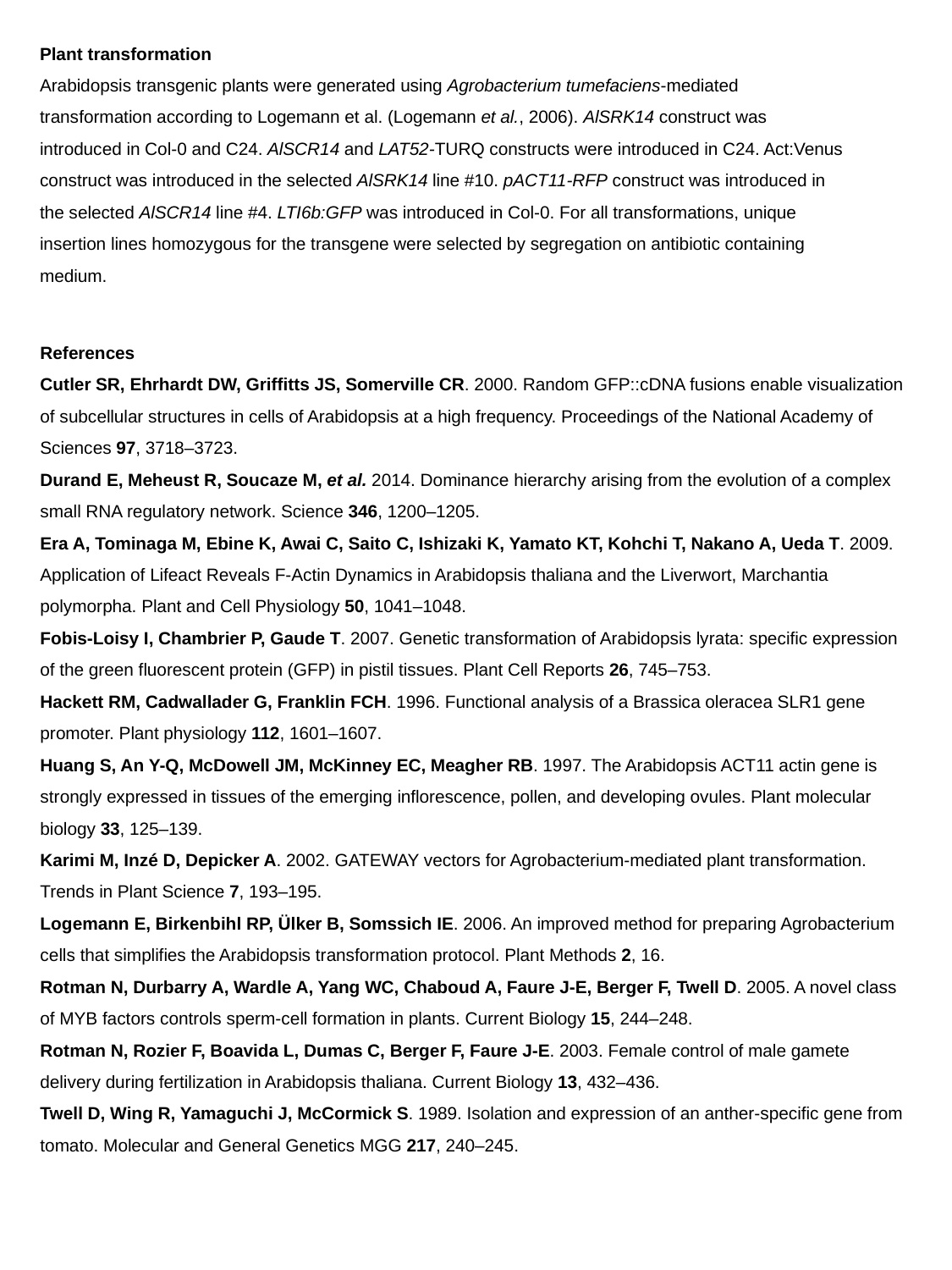

Plant transformation
Arabidopsis transgenic plants were generated using Agrobacterium tumefaciens-mediated transformation according to Logemann et al. (Logemann et al., 2006). AlSRK14 construct was
introduced in Col-0 and C24. AlSCR14 and LAT52-TURQ constructs were introduced in C24. Act:Venus construct was introduced in the selected AlSRK14 line #10. pACT11-RFP construct was introduced in the selected AlSCR14 line #4. LTI6b:GFP was introduced in Col-0. For all transformations, unique insertion lines homozygous for the transgene were selected by segregation on antibiotic containing medium.
References
Cutler SR, Ehrhardt DW, Griffitts JS, Somerville CR. 2000. Random GFP::cDNA fusions enable visualization of subcellular structures in cells of Arabidopsis at a high frequency. Proceedings of the National Academy of Sciences 97, 3718–3723.
Durand E, Meheust R, Soucaze M, et al. 2014. Dominance hierarchy arising from the evolution of a complex small RNA regulatory network. Science 346, 1200–1205.
Era A, Tominaga M, Ebine K, Awai C, Saito C, Ishizaki K, Yamato KT, Kohchi T, Nakano A, Ueda T. 2009. Application of Lifeact Reveals F-Actin Dynamics in Arabidopsis thaliana and the Liverwort, Marchantia polymorpha. Plant and Cell Physiology 50, 1041–1048.
Fobis-Loisy I, Chambrier P, Gaude T. 2007. Genetic transformation of Arabidopsis lyrata: specific expression of the green fluorescent protein (GFP) in pistil tissues. Plant Cell Reports 26, 745–753.
Hackett RM, Cadwallader G, Franklin FCH. 1996. Functional analysis of a Brassica oleracea SLR1 gene promoter. Plant physiology 112, 1601–1607.
Huang S, An Y-Q, McDowell JM, McKinney EC, Meagher RB. 1997. The Arabidopsis ACT11 actin gene is strongly expressed in tissues of the emerging inflorescence, pollen, and developing ovules. Plant molecular biology 33, 125–139.
Karimi M, Inzé D, Depicker A. 2002. GATEWAY vectors for Agrobacterium-mediated plant transformation. Trends in Plant Science 7, 193–195.
Logemann E, Birkenbihl RP, Ülker B, Somssich IE. 2006. An improved method for preparing Agrobacterium cells that simplifies the Arabidopsis transformation protocol. Plant Methods 2, 16.
Rotman N, Durbarry A, Wardle A, Yang WC, Chaboud A, Faure J-E, Berger F, Twell D. 2005. A novel class of MYB factors controls sperm-cell formation in plants. Current Biology 15, 244–248.
Rotman N, Rozier F, Boavida L, Dumas C, Berger F, Faure J-E. 2003. Female control of male gamete delivery during fertilization in Arabidopsis thaliana. Current Biology 13, 432–436.
Twell D, Wing R, Yamaguchi J, McCormick S. 1989. Isolation and expression of an anther-specific gene from tomato. Molecular and General Genetics MGG 217, 240–245.

### Slide 4
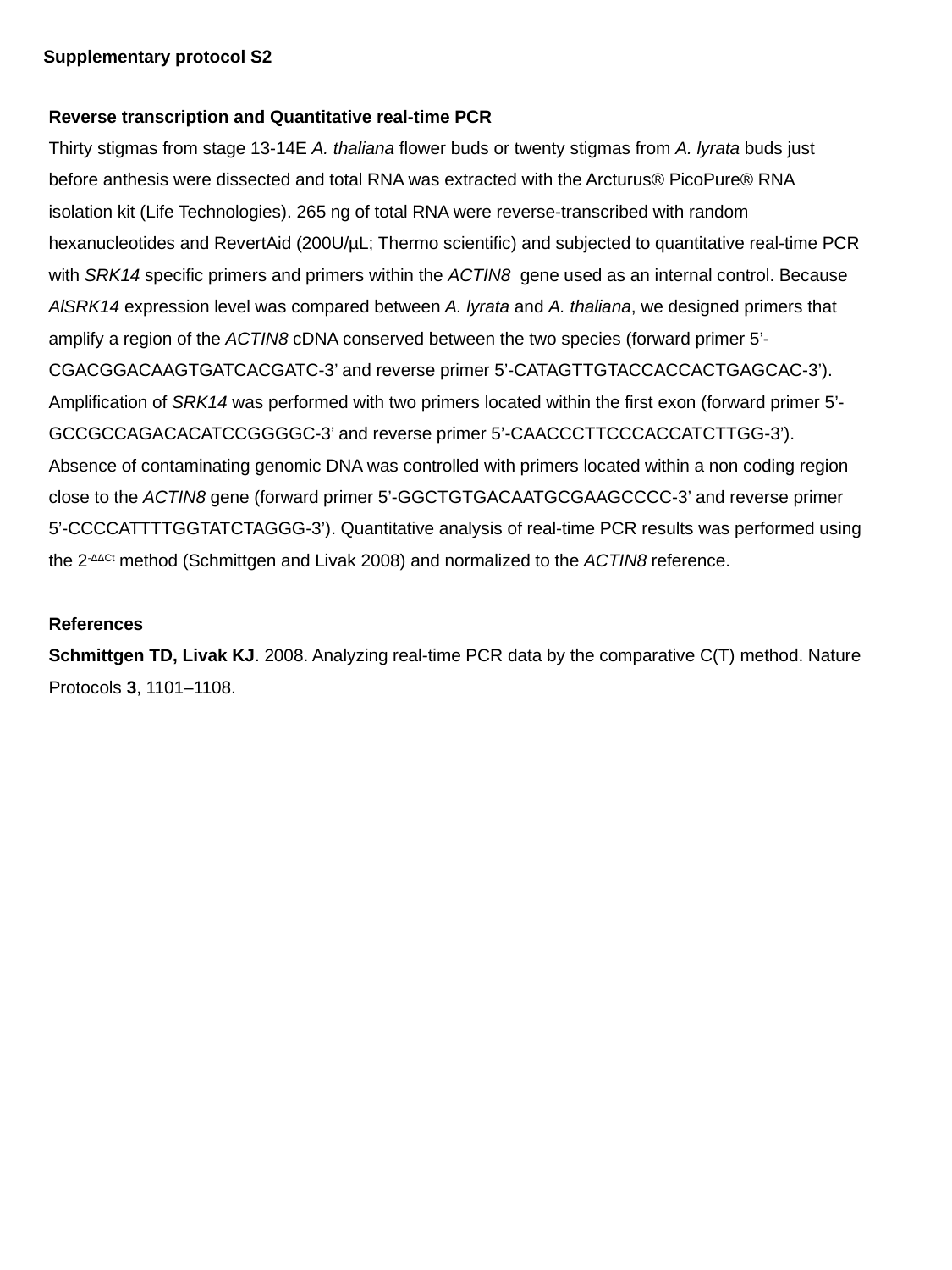

Supplementary protocol S2
Reverse transcription and Quantitative real-time PCR
Thirty stigmas from stage 13-14E A. thaliana flower buds or twenty stigmas from A. lyrata buds just before anthesis were dissected and total RNA was extracted with the Arcturus® PicoPure® RNA isolation kit (Life Technologies). 265 ng of total RNA were reverse-transcribed with random hexanucleotides and RevertAid (200U/µL; Thermo scientific) and subjected to quantitative real-time PCR with SRK14 specific primers and primers within the ACTIN8 gene used as an internal control. Because AlSRK14 expression level was compared between A. lyrata and A. thaliana, we designed primers that amplify a region of the ACTIN8 cDNA conserved between the two species (forward primer 5’-CGACGGACAAGTGATCACGATC-3’ and reverse primer 5’-CATAGTTGTACCACCACTGAGCAC-3’). Amplification of SRK14 was performed with two primers located within the first exon (forward primer 5’-GCCGCCAGACACATCCGGGGC-3’ and reverse primer 5’-CAACCCTTCCCACCATCTTGG-3’). Absence of contaminating genomic DNA was controlled with primers located within a non coding region close to the ACTIN8 gene (forward primer 5’-GGCTGTGACAATGCGAAGCCCC-3’ and reverse primer 5’-CCCCATTTTGGTATCTAGGG-3’). Quantitative analysis of real-time PCR results was performed using the 2^-ΔΔCt^ method (Schmittgen and Livak 2008) and normalized to the ACTIN8 reference.
References
Schmittgen TD, Livak KJ. 2008. Analyzing real-time PCR data by the comparative C(T) method. Nature Protocols 3, 1101–1108.

### Slide 5
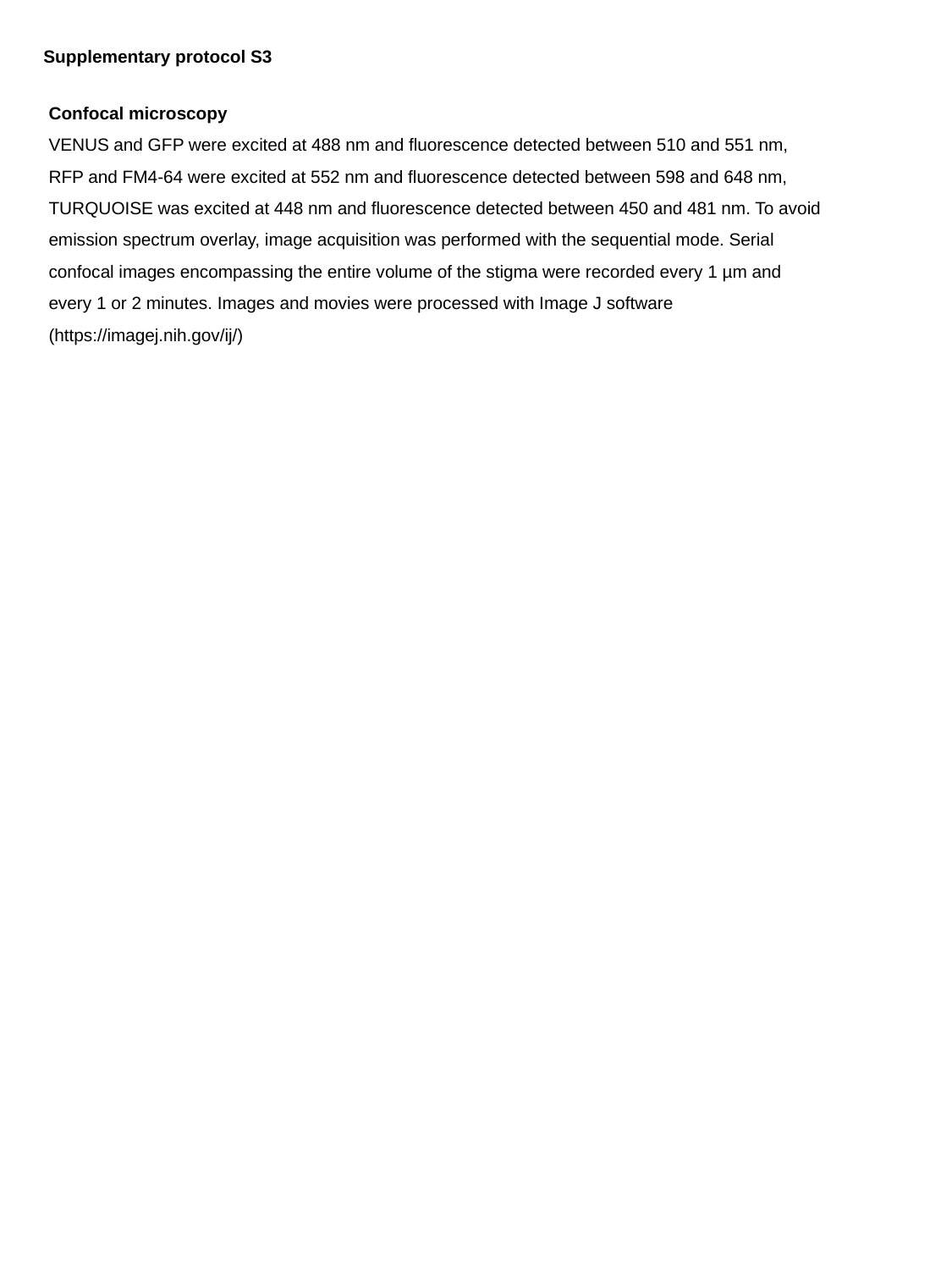

Supplementary protocol S3
Confocal microscopy
VENUS and GFP were excited at 488 nm and fluorescence detected between 510 and 551 nm, RFP and FM4-64 were excited at 552 nm and fluorescence detected between 598 and 648 nm, TURQUOISE was excited at 448 nm and fluorescence detected between 450 and 481 nm. To avoid emission spectrum overlay, image acquisition was performed with the sequential mode. Serial confocal images encompassing the entire volume of the stigma were recorded every 1 µm and every 1 or 2 minutes. Images and movies were processed with Image J software (https://imagej.nih.gov/ij/)

### Slide 6
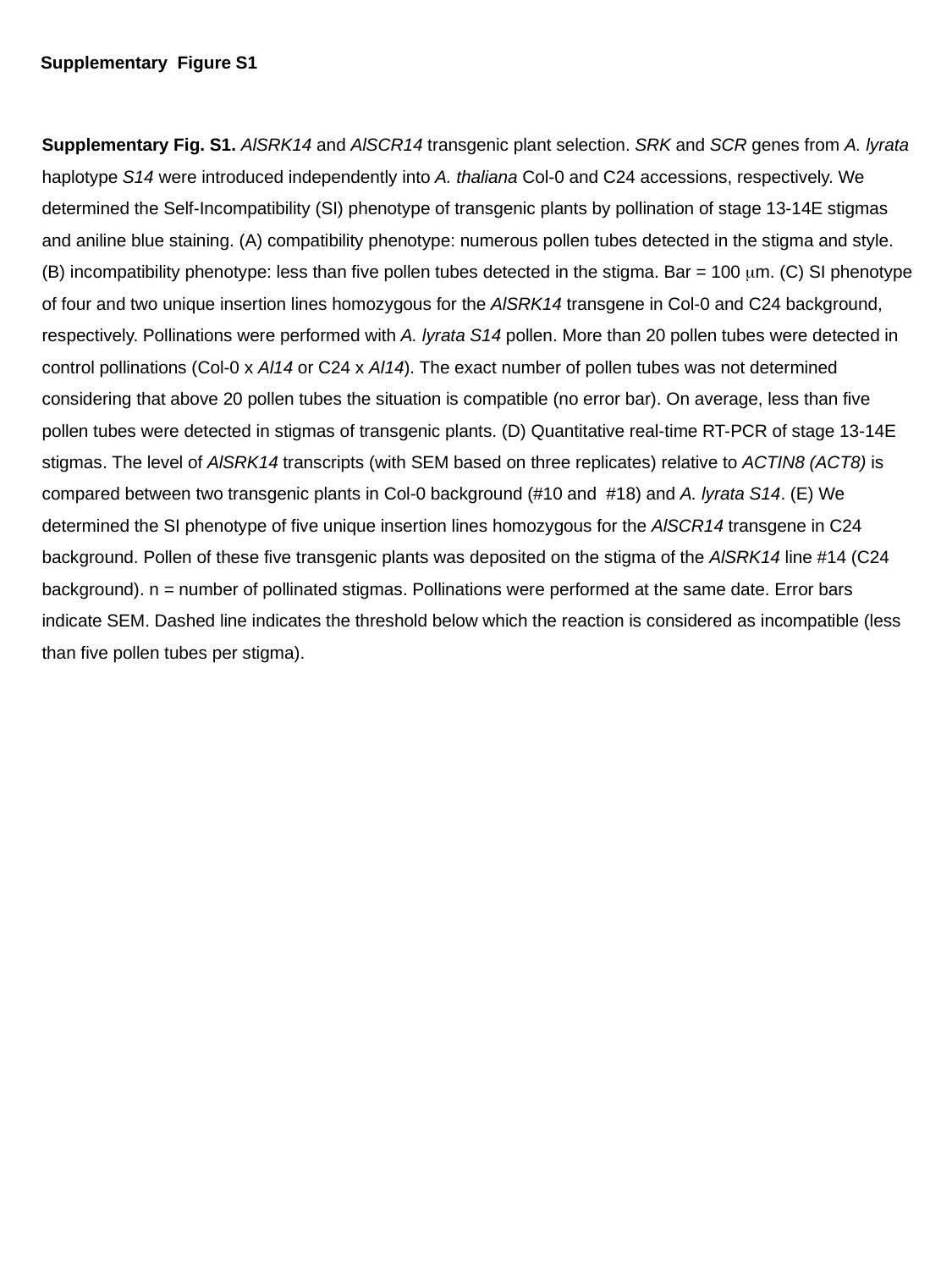

Supplementary Figure S1
Supplementary Fig. S1. AlSRK14 and AlSCR14 transgenic plant selection. SRK and SCR genes from A. lyrata haplotype S14 were introduced independently into A. thaliana Col-0 and C24 accessions, respectively. We determined the Self-Incompatibility (SI) phenotype of transgenic plants by pollination of stage 13-14E stigmas and aniline blue staining. (A) compatibility phenotype: numerous pollen tubes detected in the stigma and style. (B) incompatibility phenotype: less than five pollen tubes detected in the stigma. Bar = 100 μm. (C) SI phenotype of four and two unique insertion lines homozygous for the AlSRK14 transgene in Col-0 and C24 background, respectively. Pollinations were performed with A. lyrata S14 pollen. More than 20 pollen tubes were detected in control pollinations (Col-0 x Al14 or C24 x Al14). The exact number of pollen tubes was not determined considering that above 20 pollen tubes the situation is compatible (no error bar). On average, less than five pollen tubes were detected in stigmas of transgenic plants. (D) Quantitative real-time RT-PCR of stage 13-14E stigmas. The level of AlSRK14 transcripts (with SEM based on three replicates) relative to ACTIN8 (ACT8) is compared between two transgenic plants in Col-0 background (#10 and #18) and A. lyrata S14. (E) We determined the SI phenotype of five unique insertion lines homozygous for the AlSCR14 transgene in C24 background. Pollen of these five transgenic plants was deposited on the stigma of the AlSRK14 line #14 (C24 background). n = number of pollinated stigmas. Pollinations were performed at the same date. Error bars indicate SEM. Dashed line indicates the threshold below which the reaction is considered as incompatible (less than five pollen tubes per stigma).

### Slide 7
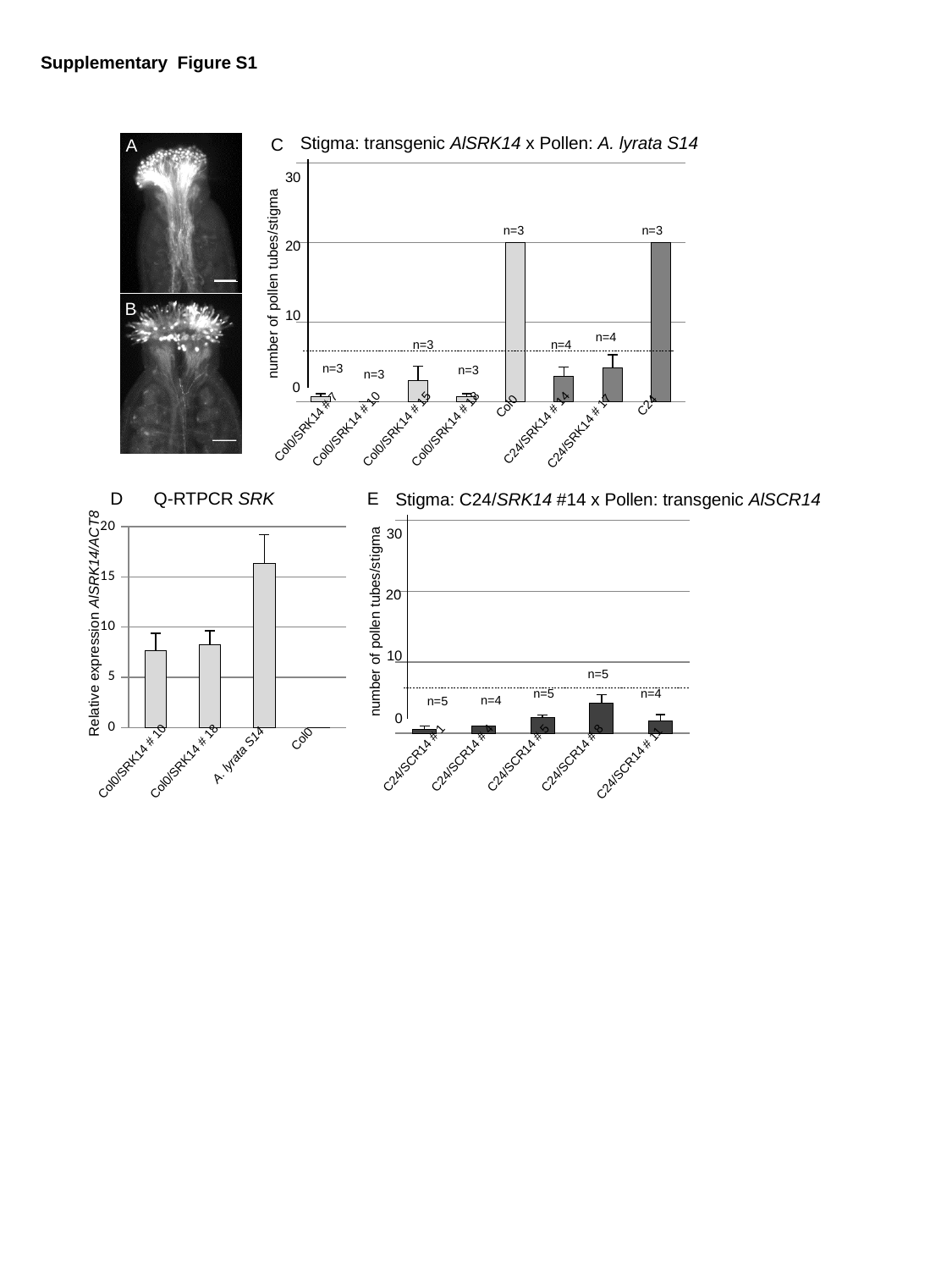

Supplementary Figure S1
Stigma: transgenic AlSRK14 x Pollen: A. lyrata S14
C
A
#### Chart
| Category | |
|---|---|30
n=3
n=3
20
number of pollen tubes/stigma
B
10
n=4
n=3
n=4
n=3
n=3
n=3
0
C24
Col0
Col0/SRK14 # 7
C24/SRK14 # 14
Col0/SRK14 # 10
Col0/SRK14 # 15
Col0/SRK14 # 18
C24/SRK14 # 17
D
Q-RTPCR SRK
E
Stigma: C24/SRK14 #14 x Pollen: transgenic AlSCR14
#### Chart
| Category |
|---|
#### Chart
| Category | |
|---|---|30
20
Relative expression AlSRK14/ACT8
number of pollen tubes/stigma
10
n=5
n=5
n=4
n=4
n=5
0
Col0
A. lyrata S14
C24/SCR14 # 1
C24/SCR14 # 4
C24/SCR14 # 5
C24/SCR14 # 8
Col0/SRK14 # 10
Col0/SRK14 # 18
C24/SCR14 # 11

### Slide 8
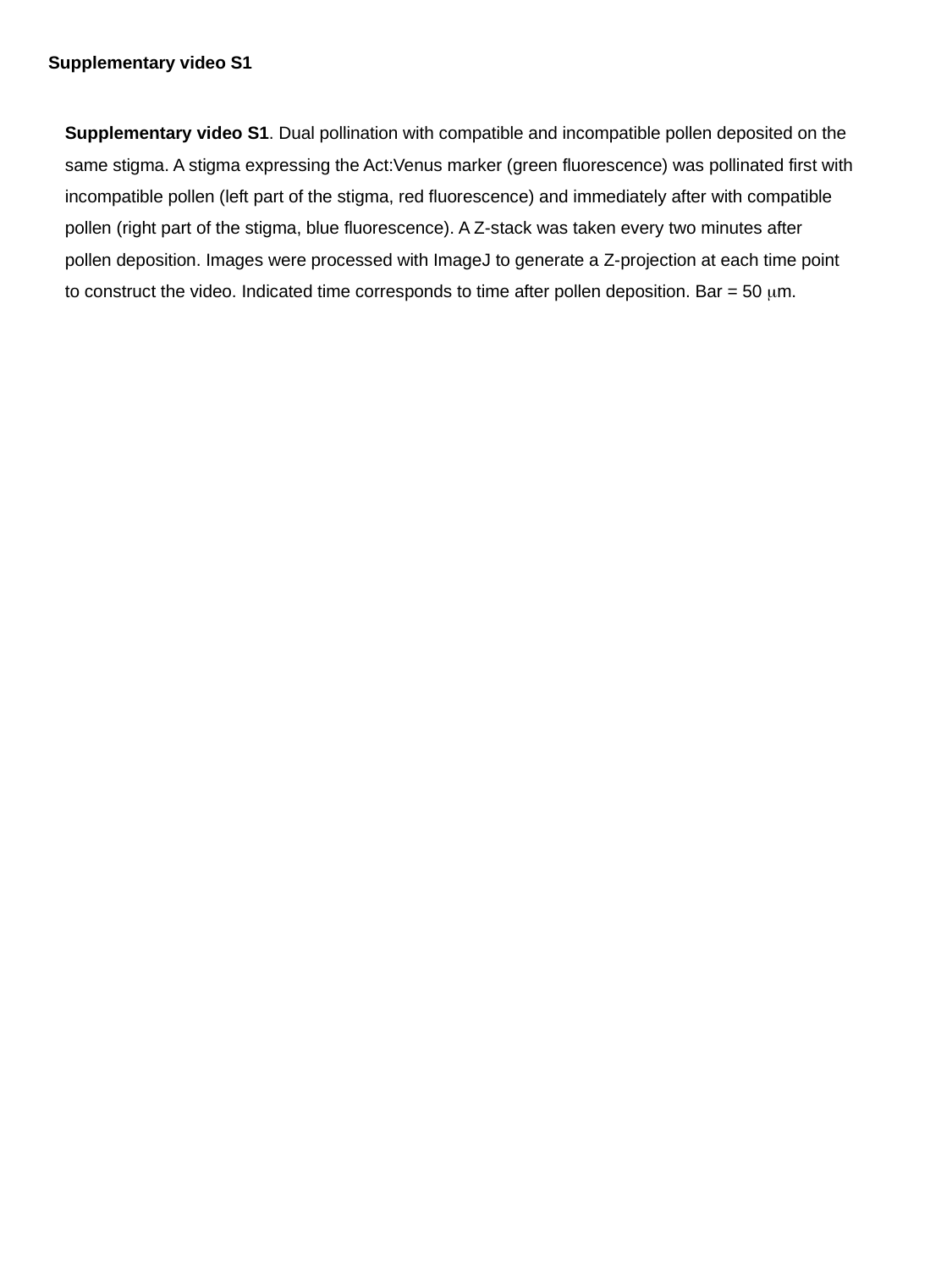

Supplementary video S1
Supplementary video S1. Dual pollination with compatible and incompatible pollen deposited on the same stigma. A stigma expressing the Act:Venus marker (green fluorescence) was pollinated first with incompatible pollen (left part of the stigma, red fluorescence) and immediately after with compatible pollen (right part of the stigma, blue fluorescence). A Z-stack was taken every two minutes after pollen deposition. Images were processed with ImageJ to generate a Z-projection at each time point to construct the video. Indicated time corresponds to time after pollen deposition. Bar = 50 μm.

### Slide 9
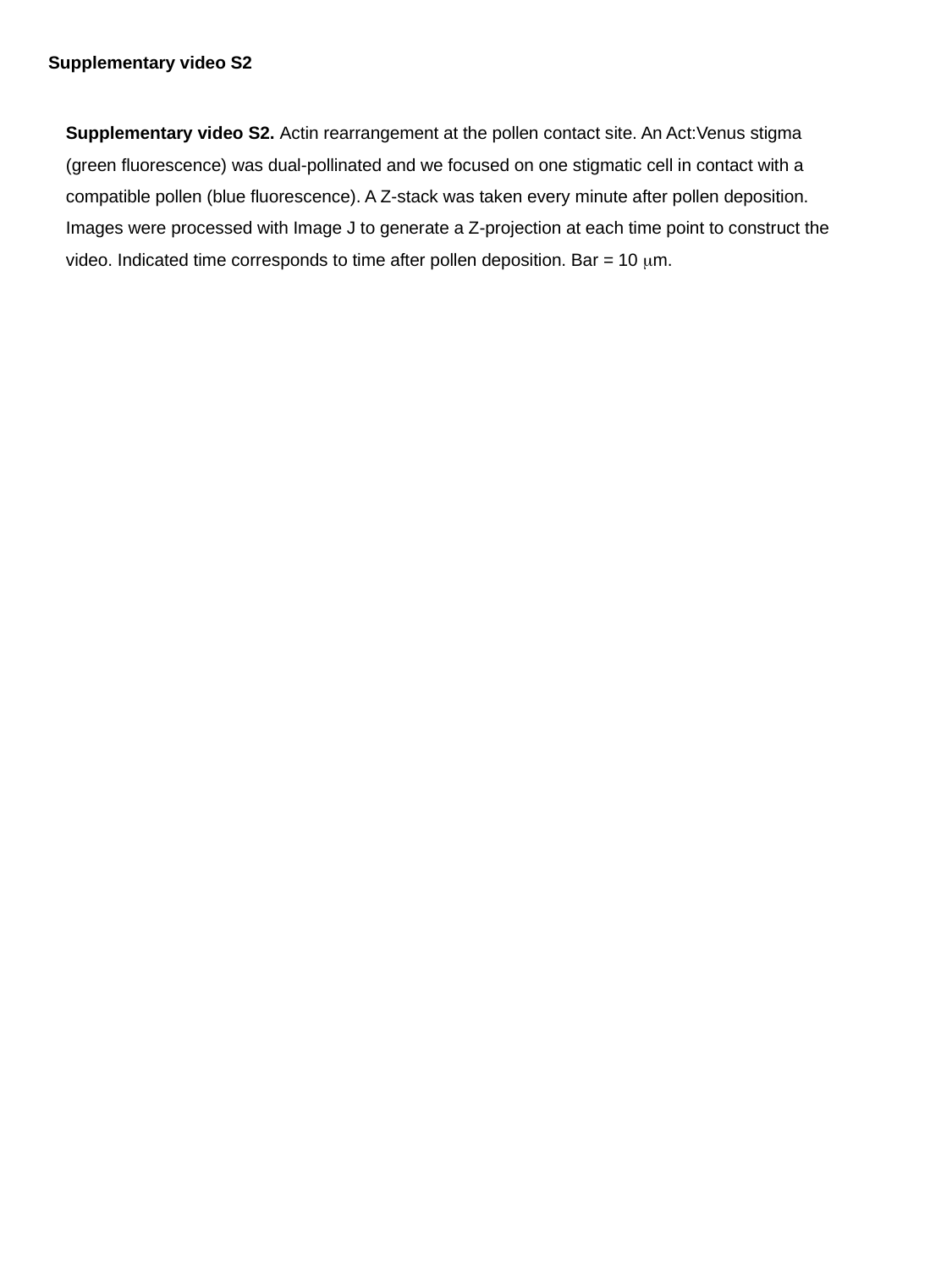

Supplementary video S2
Supplementary video S2. Actin rearrangement at the pollen contact site. An Act:Venus stigma (green fluorescence) was dual-pollinated and we focused on one stigmatic cell in contact with a compatible pollen (blue fluorescence). A Z-stack was taken every minute after pollen deposition. Images were processed with Image J to generate a Z-projection at each time point to construct the video. Indicated time corresponds to time after pollen deposition. Bar = 10 μm.

### Slide 10
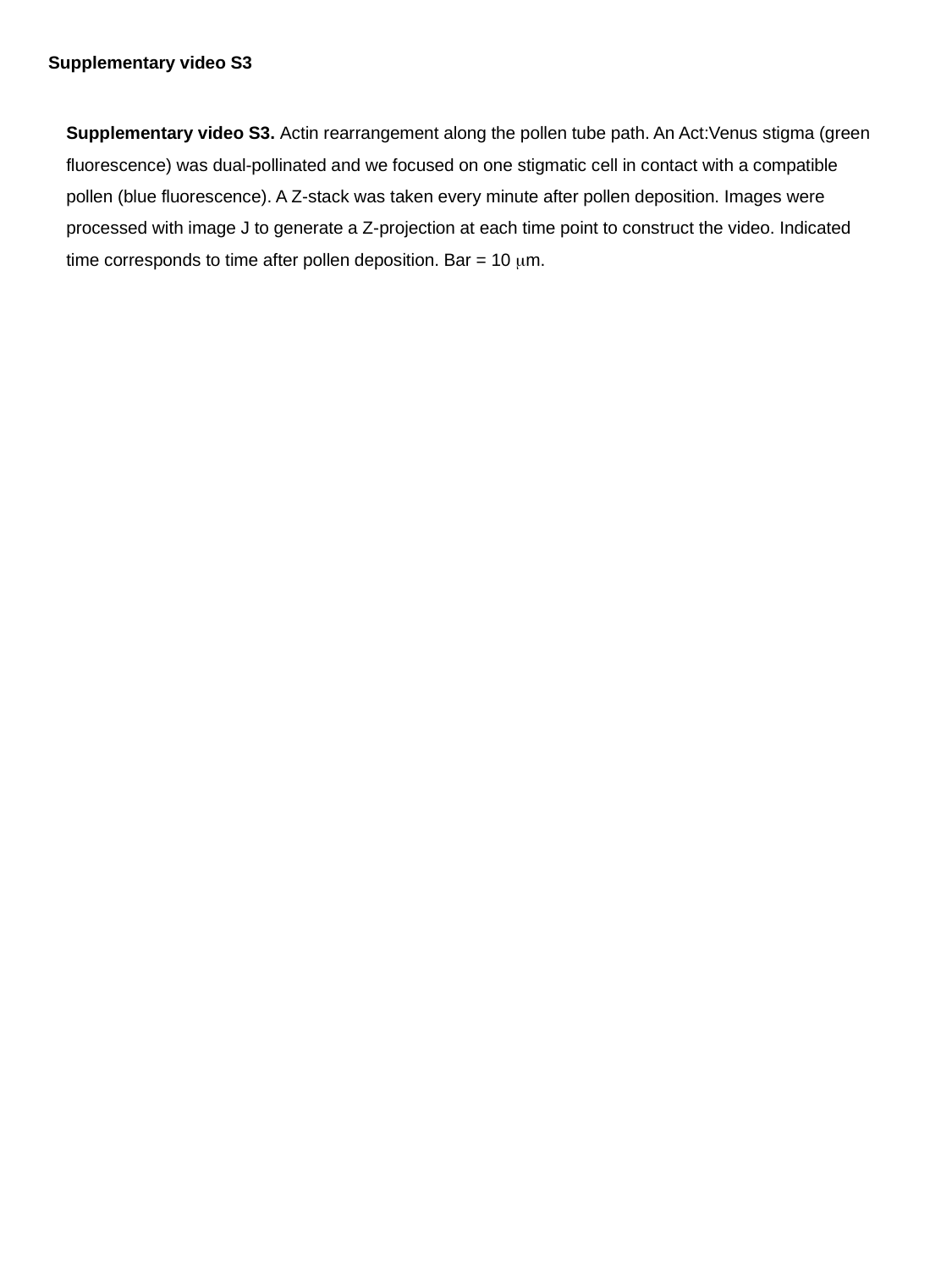

Supplementary video S3
Supplementary video S3. Actin rearrangement along the pollen tube path. An Act:Venus stigma (green fluorescence) was dual-pollinated and we focused on one stigmatic cell in contact with a compatible pollen (blue fluorescence). A Z-stack was taken every minute after pollen deposition. Images were processed with image J to generate a Z-projection at each time point to construct the video. Indicated time corresponds to time after pollen deposition. Bar = 10 μm.

### Slide 11
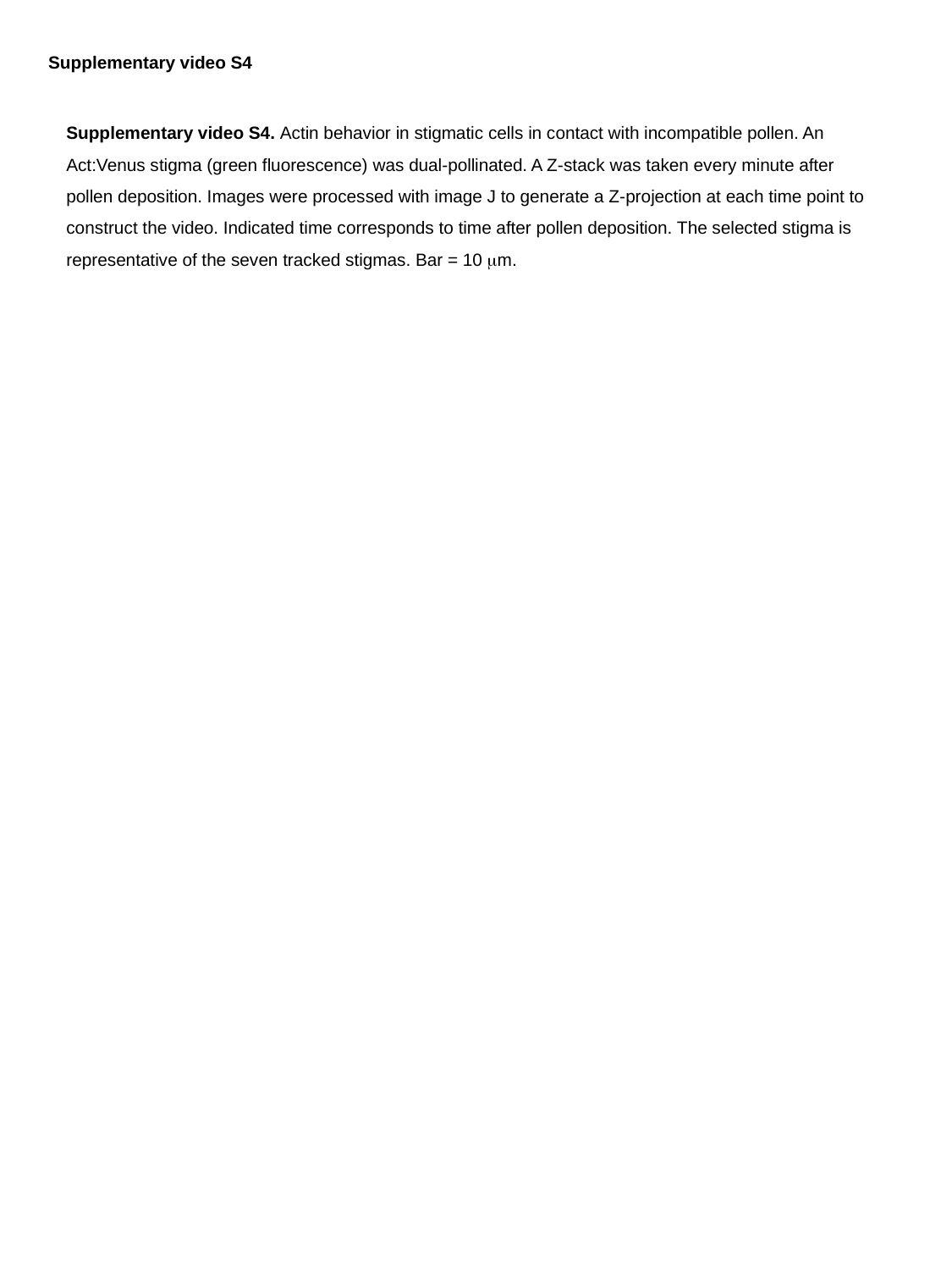

Supplementary video S4
Supplementary video S4. Actin behavior in stigmatic cells in contact with incompatible pollen. An Act:Venus stigma (green fluorescence) was dual-pollinated. A Z-stack was taken every minute after pollen deposition. Images were processed with image J to generate a Z-projection at each time point to construct the video. Indicated time corresponds to time after pollen deposition. The selected stigma is representative of the seven tracked stigmas. Bar = 10 μm.

### Slide 12
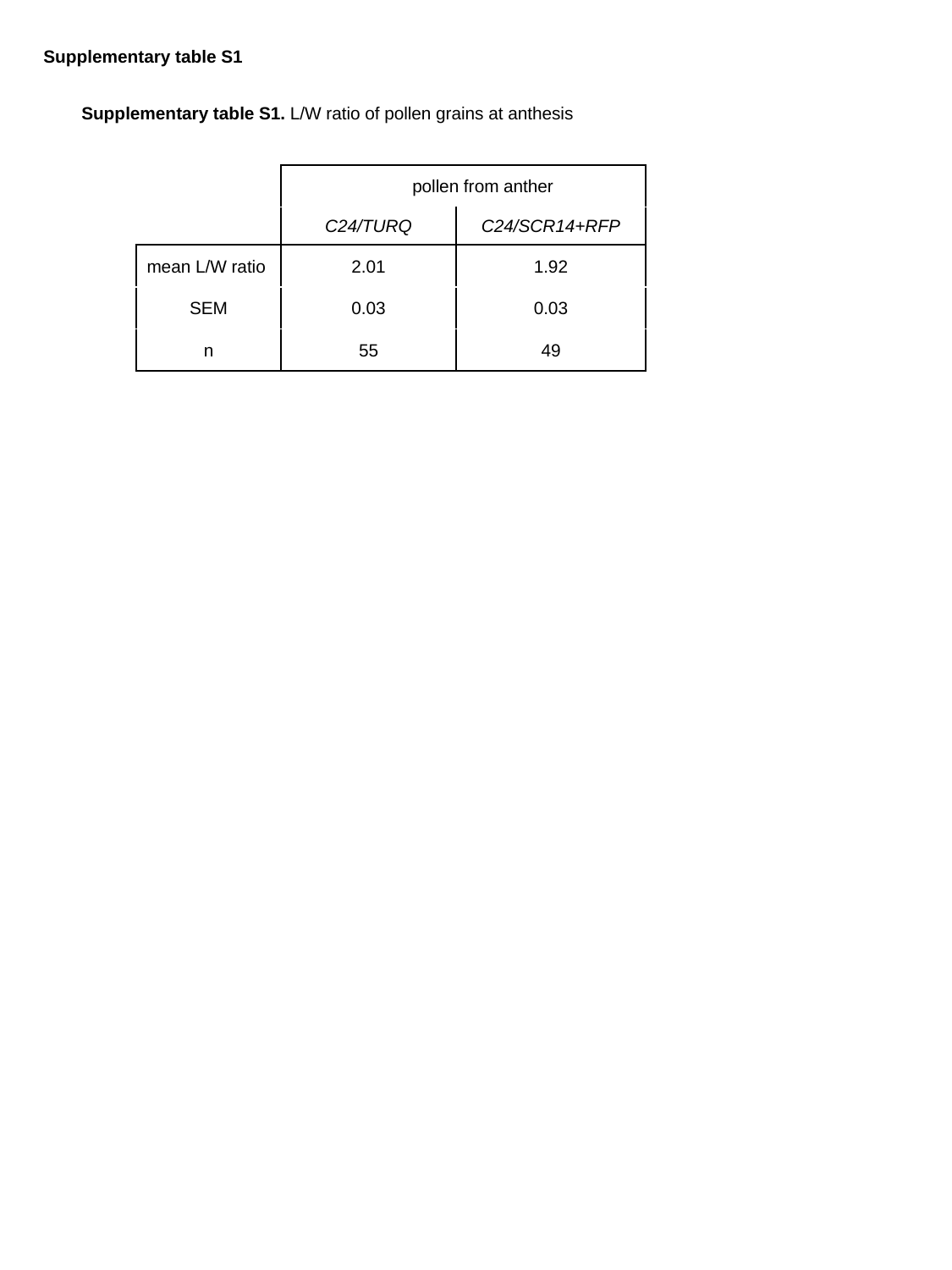

Supplementary table S1
Supplementary table S1. L/W ratio of pollen grains at anthesis
| | pollen from anther | |
| --- | --- | --- |
| | C24/TURQ | C24/SCR14+RFP |
| mean L/W ratio | 2.01 | 1.92 |
| SEM | 0.03 | 0.03 |
| n | 55 | 49 |

### Slide 13
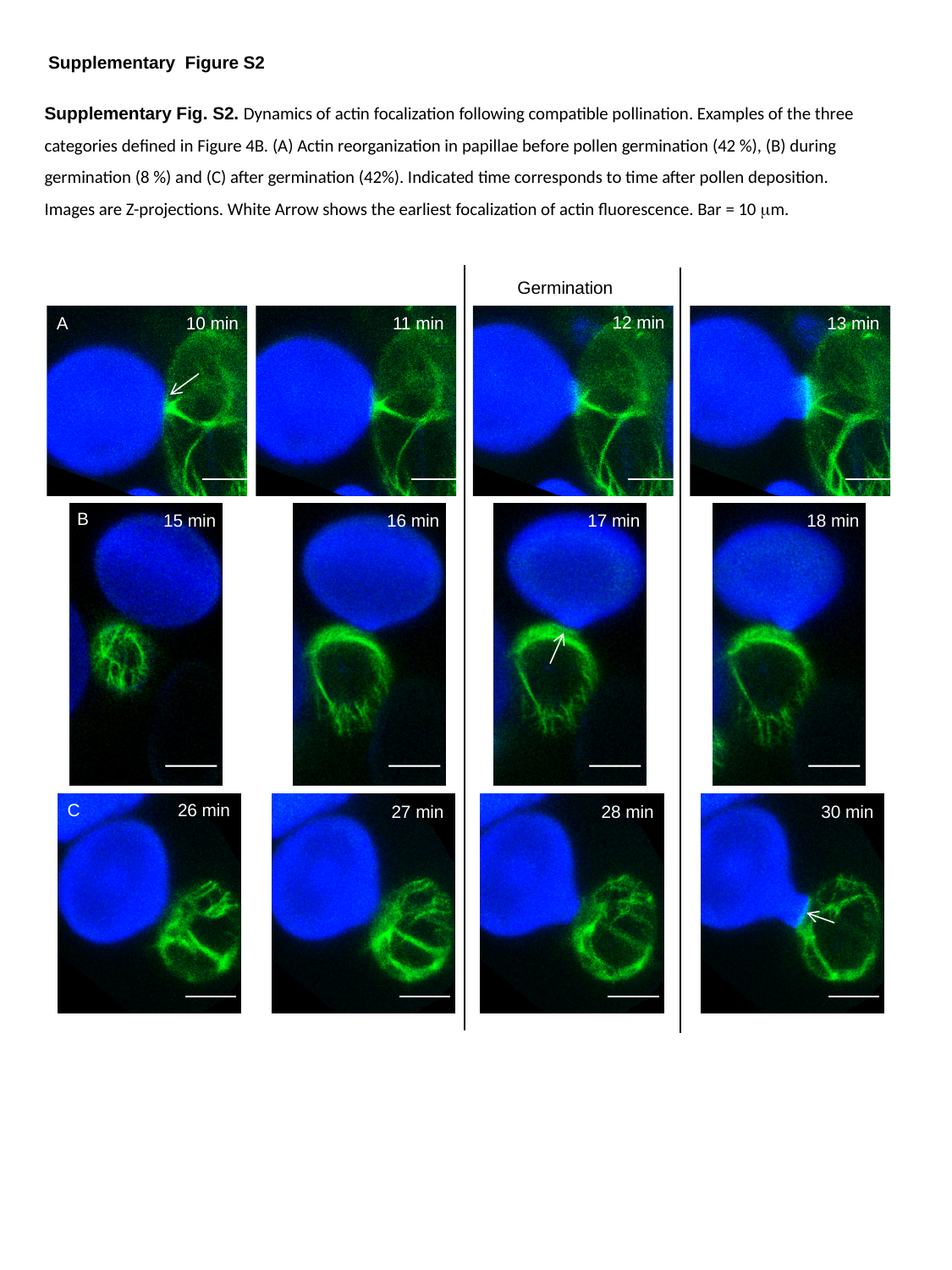

Supplementary Figure S2
Supplementary Fig. S2. Dynamics of actin focalization following compatible pollination. Examples of the three categories defined in Figure 4B. (A) Actin reorganization in papillae before pollen germination (42 %), (B) during germination (8 %) and (C) after germination (42%). Indicated time corresponds to time after pollen deposition. Images are Z-projections. White Arrow shows the earliest focalization of actin fluorescence. Bar = 10 μm.
Germination
12 min
A
10 min
11 min
13 min
B
18 min
15 min
16 min
17 min
C
26 min
28 min
30 min
27 min

### Slide 14
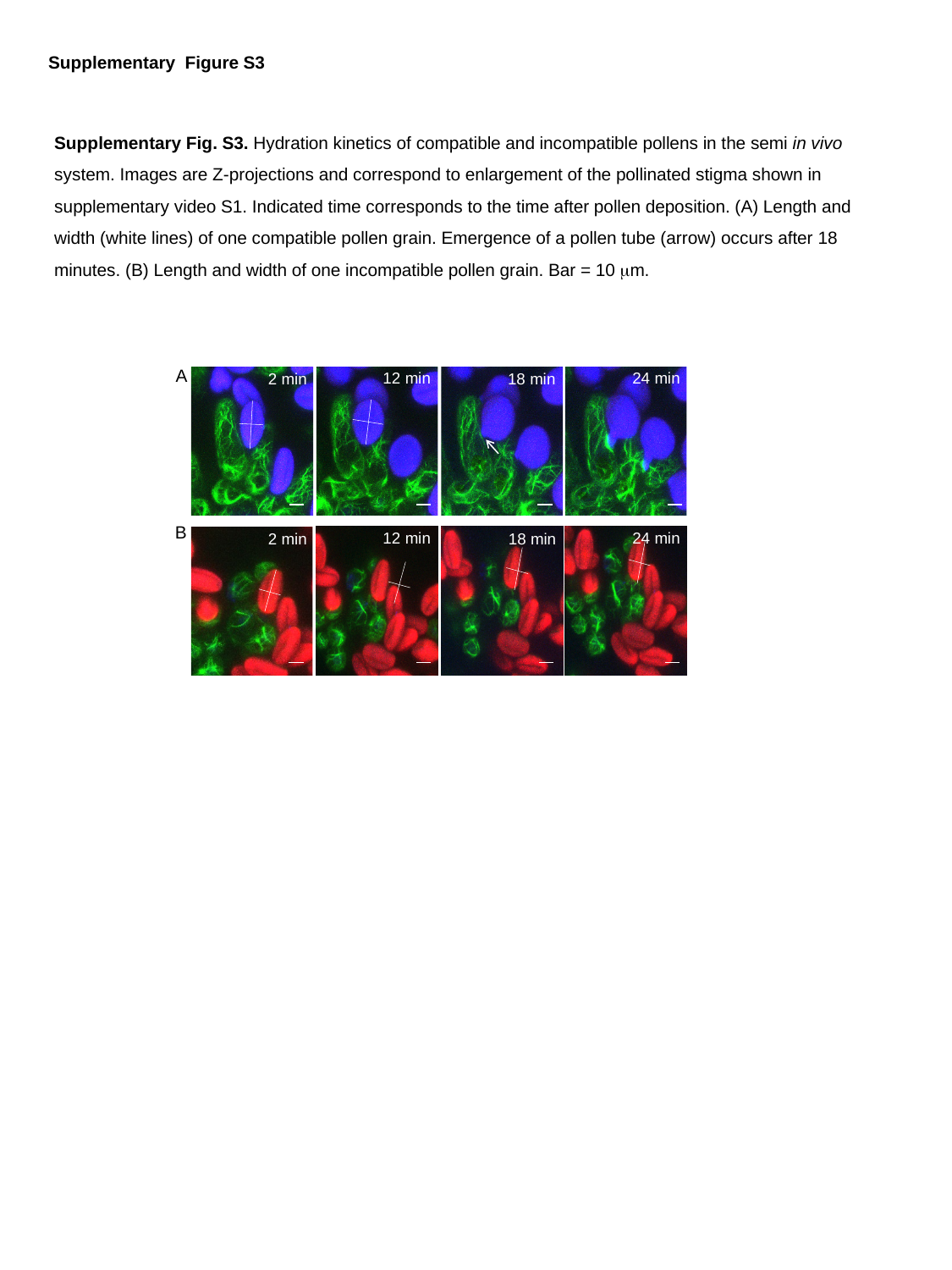

Supplementary Figure S3
Supplementary Fig. S3. Hydration kinetics of compatible and incompatible pollens in the semi in vivo system. Images are Z-projections and correspond to enlargement of the pollinated stigma shown in supplementary video S1. Indicated time corresponds to the time after pollen deposition. (A) Length and width (white lines) of one compatible pollen grain. Emergence of a pollen tube (arrow) occurs after 18 minutes. (B) Length and width of one incompatible pollen grain. Bar = 10 μm.
A
12 min
24 min
2 min
18 min
B
12 min
24 min
2 min
18 min

### Slide 15
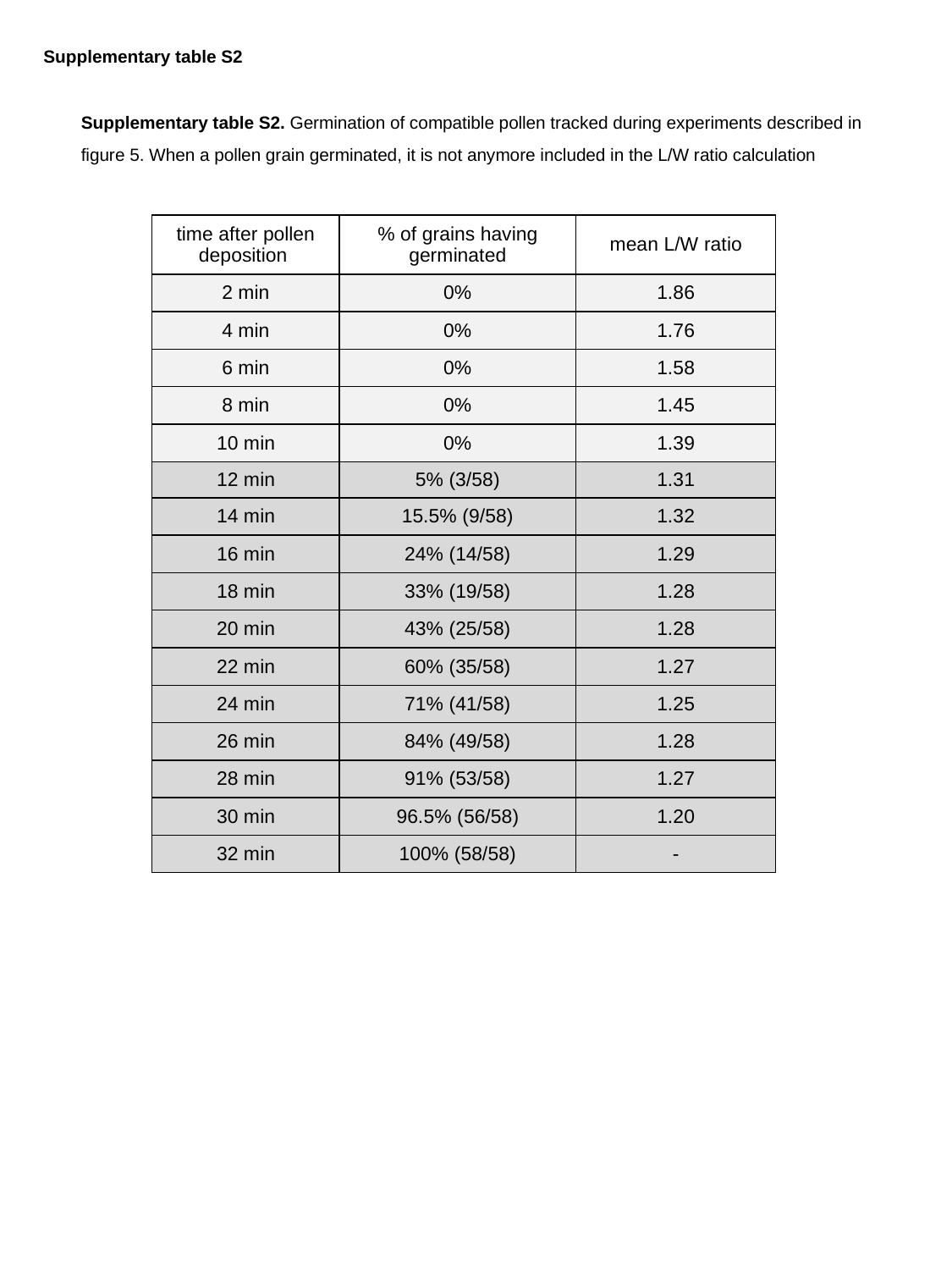

Supplementary table S2
Supplementary table S2. Germination of compatible pollen tracked during experiments described in figure 5. When a pollen grain germinated, it is not anymore included in the L/W ratio calculation
| time after pollen deposition | % of grains having germinated | mean L/W ratio |
| --- | --- | --- |
| 2 min | 0% | 1.86 |
| 4 min | 0% | 1.76 |
| 6 min | 0% | 1.58 |
| 8 min | 0% | 1.45 |
| 10 min | 0% | 1.39 |
| 12 min | 5% (3/58) | 1.31 |
| 14 min | 15.5% (9/58) | 1.32 |
| 16 min | 24% (14/58) | 1.29 |
| 18 min | 33% (19/58) | 1.28 |
| 20 min | 43% (25/58) | 1.28 |
| 22 min | 60% (35/58) | 1.27 |
| 24 min | 71% (41/58) | 1.25 |
| 26 min | 84% (49/58) | 1.28 |
| 28 min | 91% (53/58) | 1.27 |
| 30 min | 96.5% (56/58) | 1.20 |
| 32 min | 100% (58/58) | - |

### Slide 16
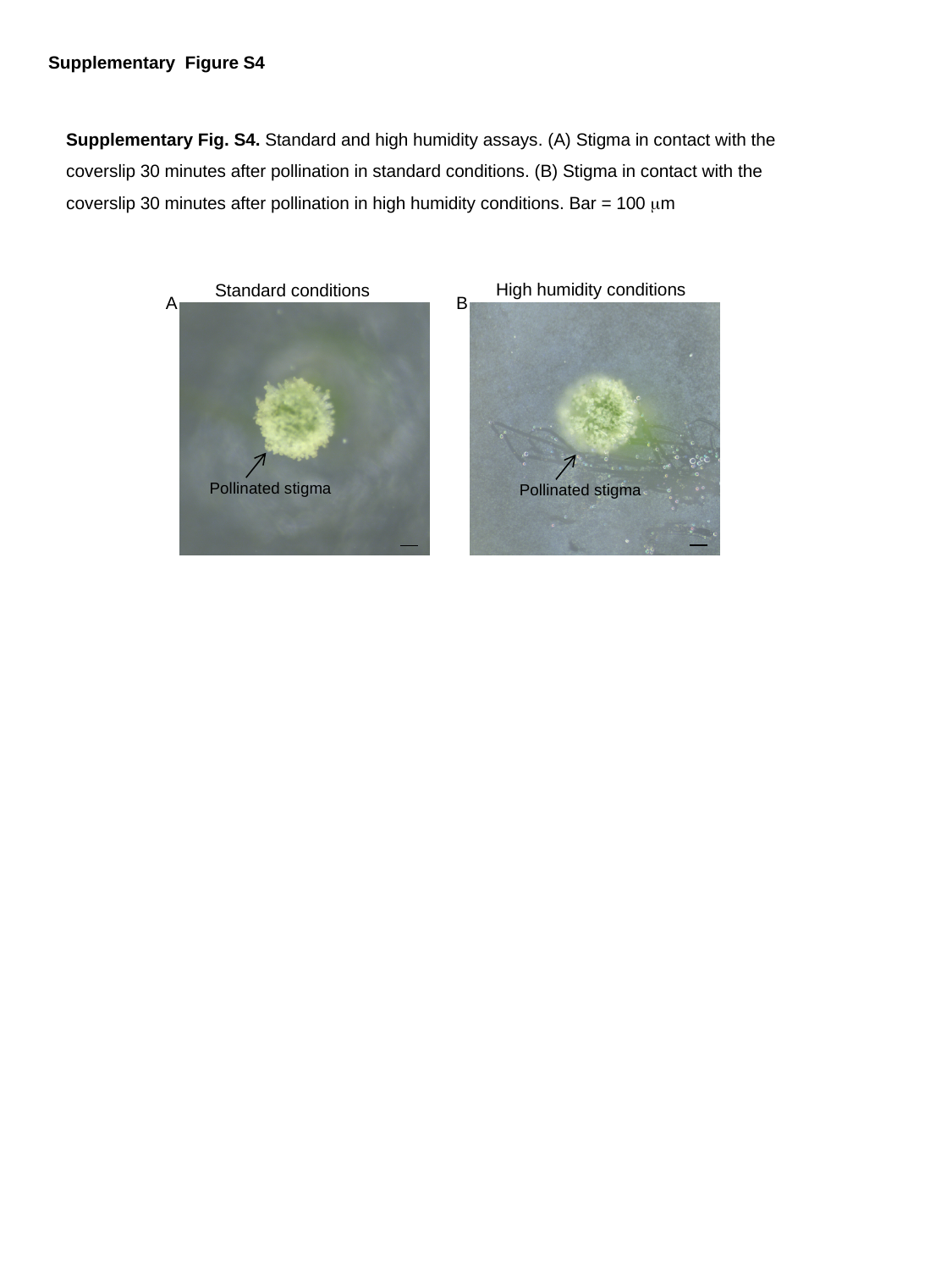

Supplementary Figure S4
Supplementary Fig. S4. Standard and high humidity assays. (A) Stigma in contact with the coverslip 30 minutes after pollination in standard conditions. (B) Stigma in contact with the coverslip 30 minutes after pollination in high humidity conditions. Bar = 100 μm
High humidity conditions
Standard conditions
B
A
Pollinated stigma
Pollinated stigma

### Slide 17
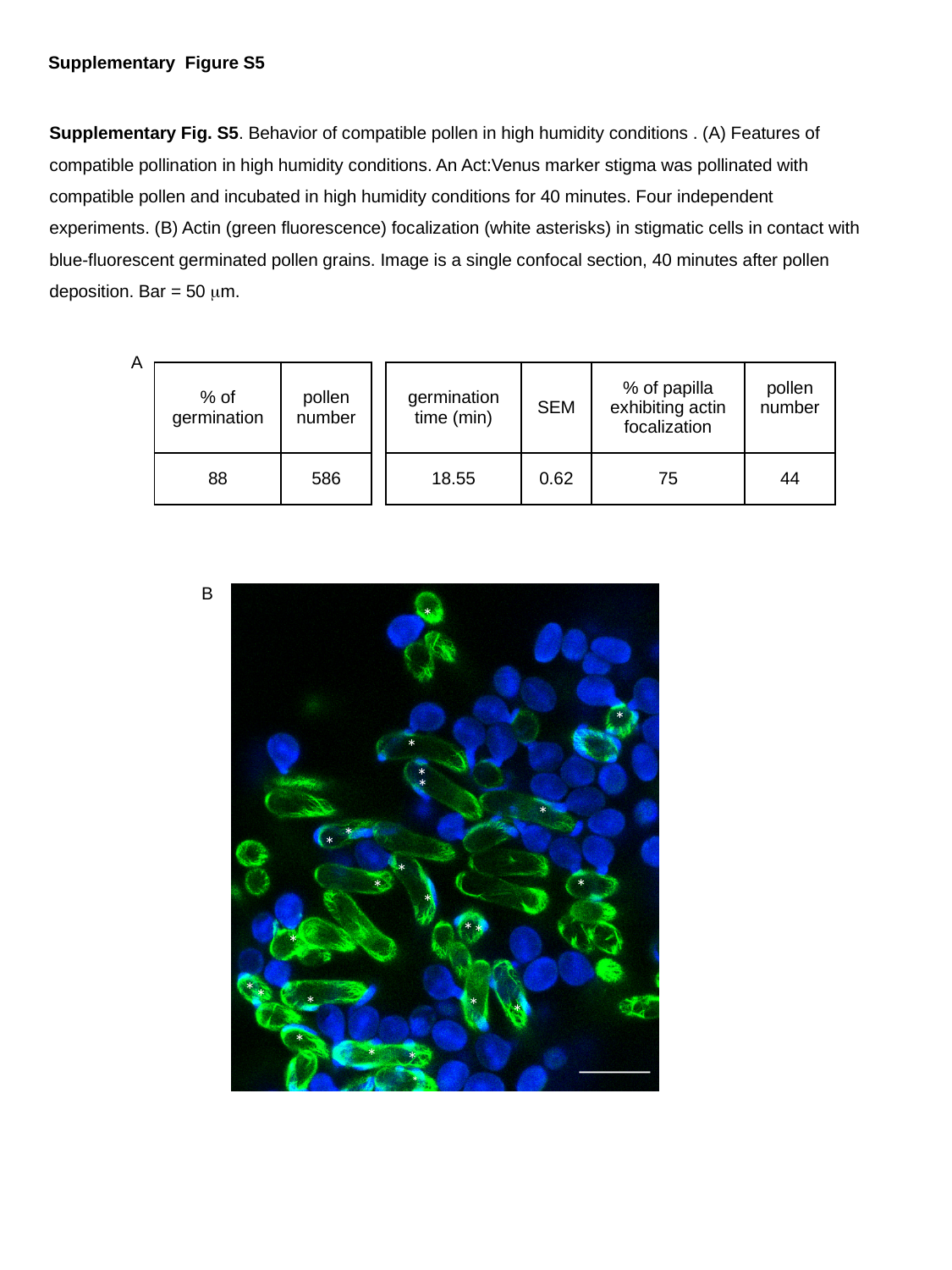

Supplementary Figure S5
Supplementary Fig. S5. Behavior of compatible pollen in high humidity conditions . (A) Features of compatible pollination in high humidity conditions. An Act:Venus marker stigma was pollinated with compatible pollen and incubated in high humidity conditions for 40 minutes. Four independent experiments. (B) Actin (green fluorescence) focalization (white asterisks) in stigmatic cells in contact with blue-fluorescent germinated pollen grains. Image is a single confocal section, 40 minutes after pollen deposition. Bar = 50 μm.
A
| % of germination | pollen number |
| --- | --- |
| 88 | 586 |
| germination time (min) | SEM | % of papilla exhibiting actin focalization | pollen number |
| --- | --- | --- | --- |
| 18.55 | 0.62 | 75 | 44 |
B
*
*
*
*
*
*
*
*
*
*
*
*
*
*
*
*
*
*
*
*
*
*
*
*
*

### Slide 18
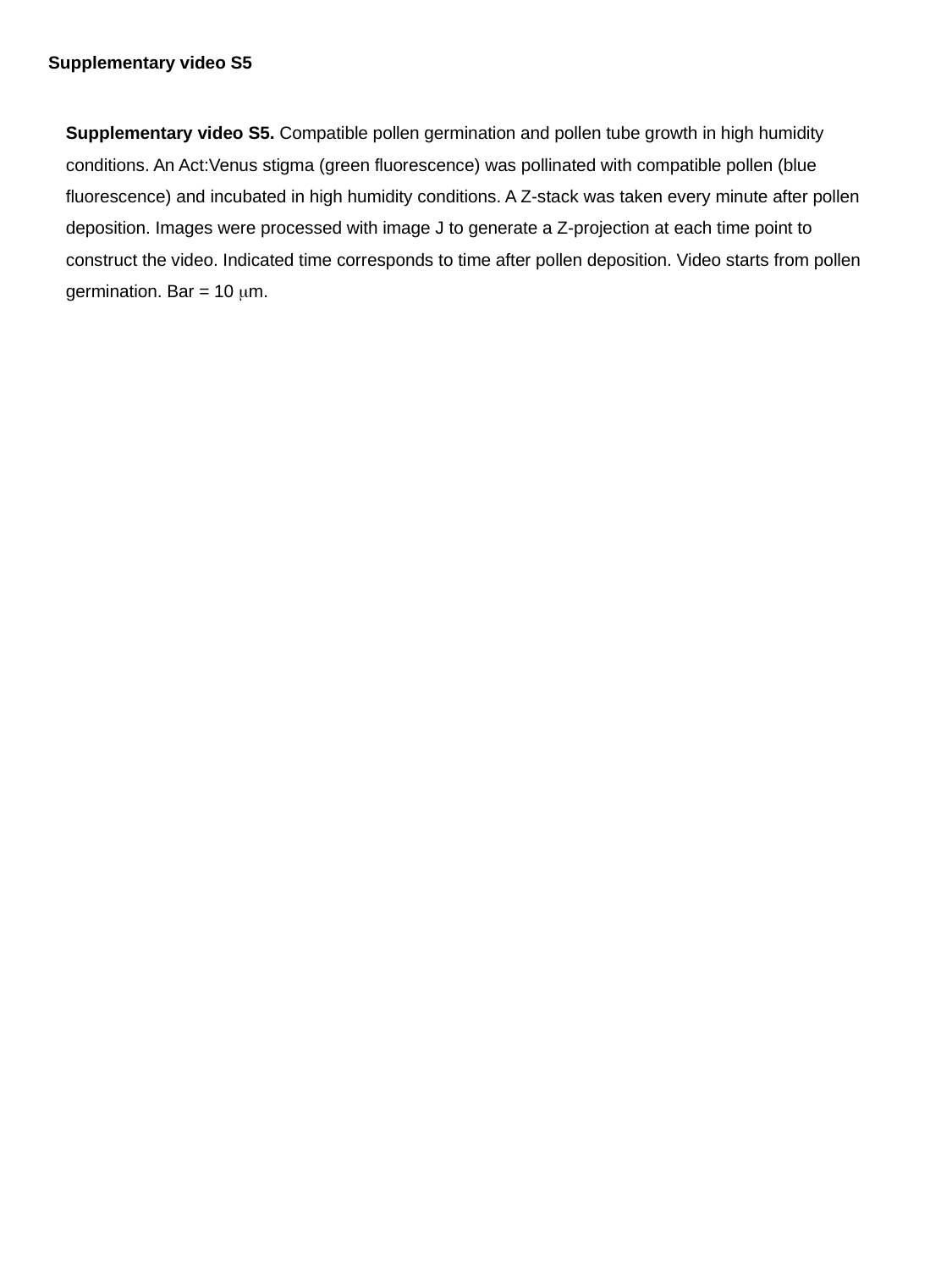

Supplementary video S5
Supplementary video S5. Compatible pollen germination and pollen tube growth in high humidity conditions. An Act:Venus stigma (green fluorescence) was pollinated with compatible pollen (blue fluorescence) and incubated in high humidity conditions. A Z-stack was taken every minute after pollen deposition. Images were processed with image J to generate a Z-projection at each time point to construct the video. Indicated time corresponds to time after pollen deposition. Video starts from pollen germination. Bar = 10 μm.

### Slide 19
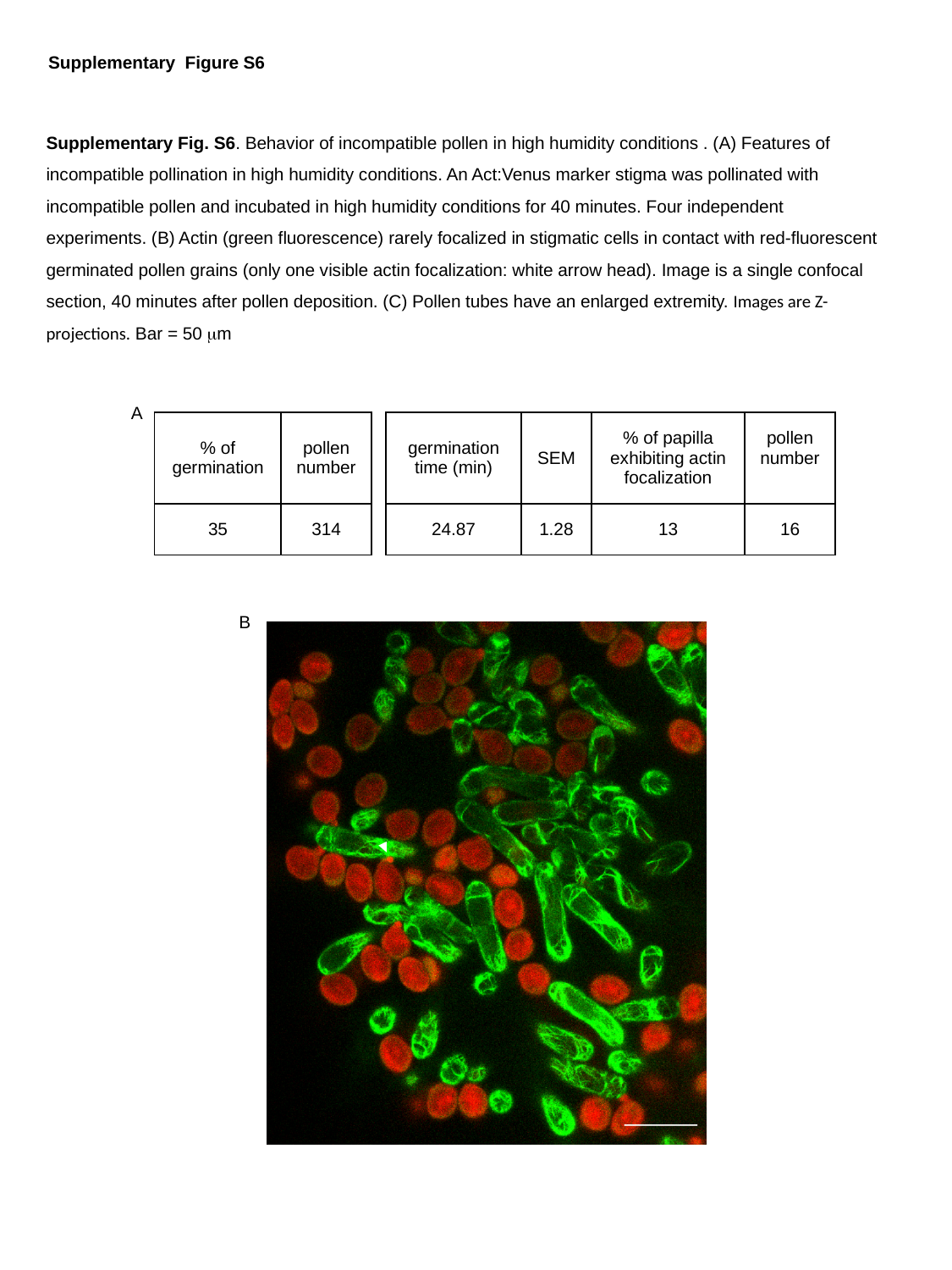

Supplementary Figure S6
Supplementary Fig. S6. Behavior of incompatible pollen in high humidity conditions . (A) Features of incompatible pollination in high humidity conditions. An Act:Venus marker stigma was pollinated with incompatible pollen and incubated in high humidity conditions for 40 minutes. Four independent experiments. (B) Actin (green fluorescence) rarely focalized in stigmatic cells in contact with red-fluorescent germinated pollen grains (only one visible actin focalization: white arrow head). Image is a single confocal section, 40 minutes after pollen deposition. (C) Pollen tubes have an enlarged extremity. Images are Z-projections. Bar = 50 μm
A
| % of germination | pollen number |
| --- | --- |
| 35 | 314 |
| germination time (min) | SEM | % of papilla exhibiting actin focalization | pollen number |
| --- | --- | --- | --- |
| 24.87 | 1.28 | 13 | 16 |
B

### Slide 20
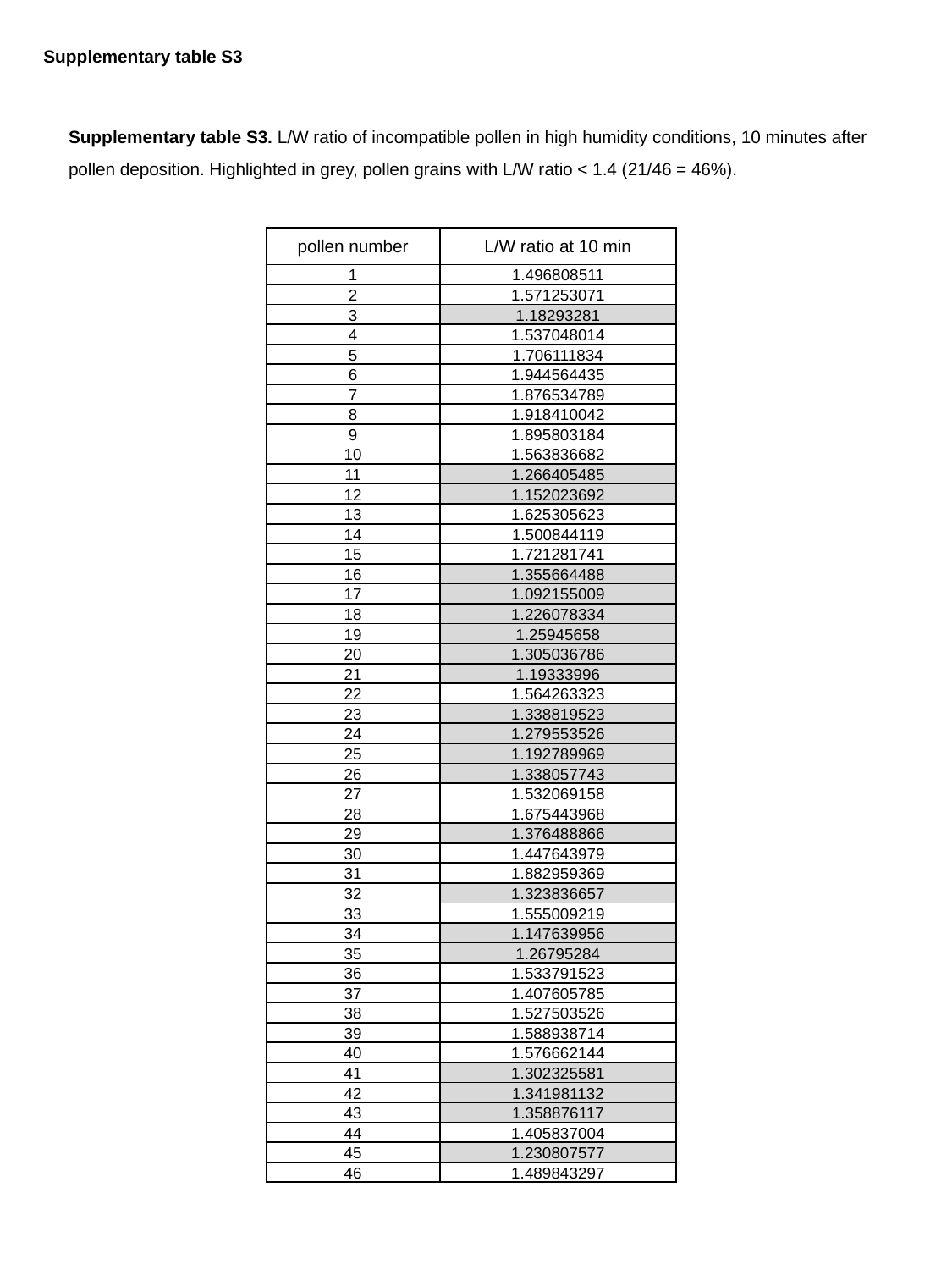

Supplementary table S3
Supplementary table S3. L/W ratio of incompatible pollen in high humidity conditions, 10 minutes after pollen deposition. Highlighted in grey, pollen grains with L/W ratio < 1.4 (21/46 = 46%).
| pollen number | L/W ratio at 10 min |
| --- | --- |
| 1 | 1.496808511 |
| 2 | 1.571253071 |
| 3 | 1.18293281 |
| 4 | 1.537048014 |
| 5 | 1.706111834 |
| 6 | 1.944564435 |
| 7 | 1.876534789 |
| 8 | 1.918410042 |
| 9 | 1.895803184 |
| 10 | 1.563836682 |
| 11 | 1.266405485 |
| 12 | 1.152023692 |
| 13 | 1.625305623 |
| 14 | 1.500844119 |
| 15 | 1.721281741 |
| 16 | 1.355664488 |
| 17 | 1.092155009 |
| 18 | 1.226078334 |
| 19 | 1.25945658 |
| 20 | 1.305036786 |
| 21 | 1.19333996 |
| 22 | 1.564263323 |
| 23 | 1.338819523 |
| 24 | 1.279553526 |
| 25 | 1.192789969 |
| 26 | 1.338057743 |
| 27 | 1.532069158 |
| 28 | 1.675443968 |
| 29 | 1.376488866 |
| 30 | 1.447643979 |
| 31 | 1.882959369 |
| 32 | 1.323836657 |
| 33 | 1.555009219 |
| 34 | 1.147639956 |
| 35 | 1.26795284 |
| 36 | 1.533791523 |
| 37 | 1.407605785 |
| 38 | 1.527503526 |
| 39 | 1.588938714 |
| 40 | 1.576662144 |
| 41 | 1.302325581 |
| 42 | 1.341981132 |
| 43 | 1.358876117 |
| 44 | 1.405837004 |
| 45 | 1.230807577 |
| 46 | 1.489843297 |

### Slide 21
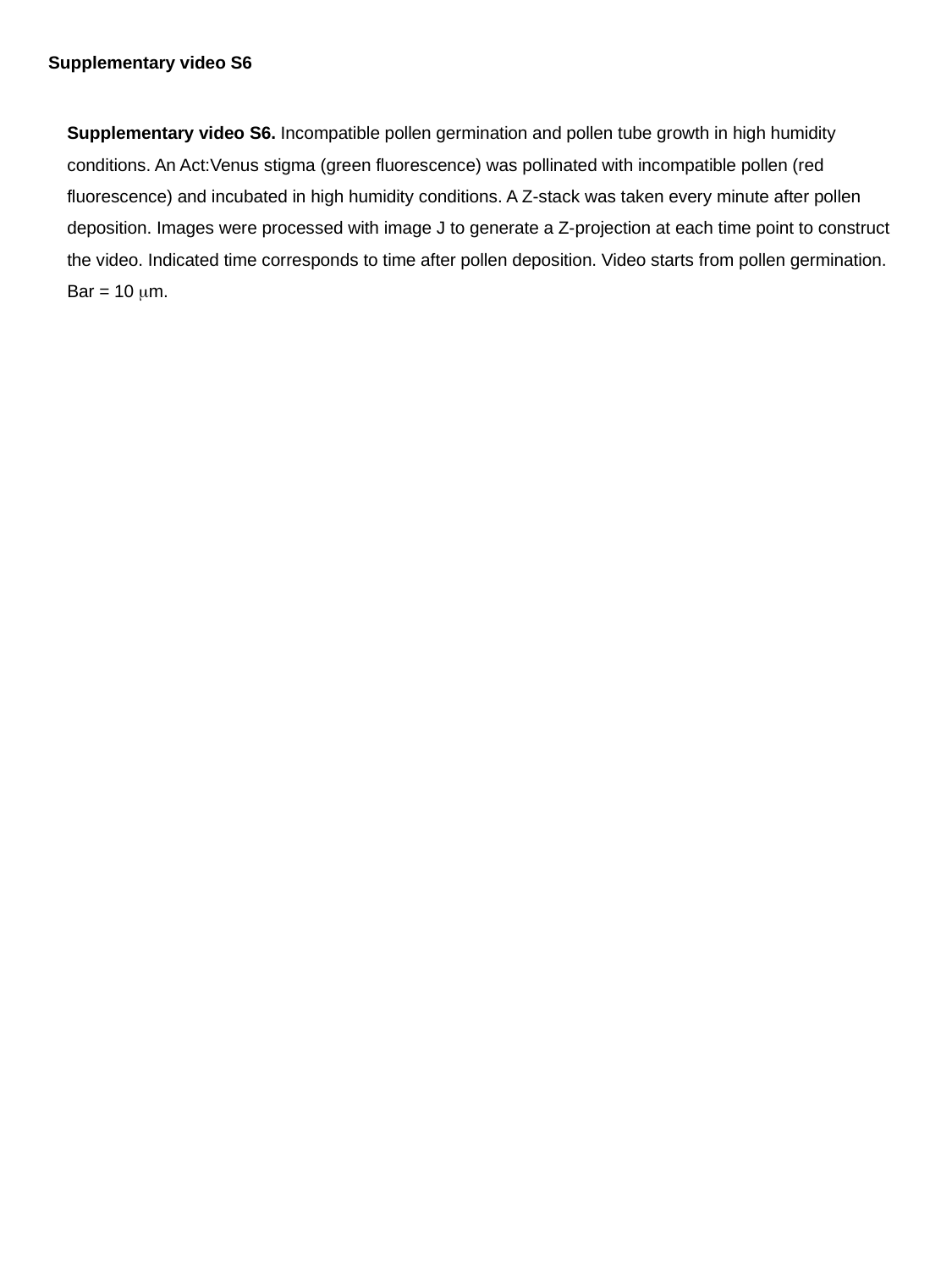

Supplementary video S6
Supplementary video S6. Incompatible pollen germination and pollen tube growth in high humidity conditions. An Act:Venus stigma (green fluorescence) was pollinated with incompatible pollen (red fluorescence) and incubated in high humidity conditions. A Z-stack was taken every minute after pollen deposition. Images were processed with image J to generate a Z-projection at each time point to construct the video. Indicated time corresponds to time after pollen deposition. Video starts from pollen germination. Bar = 10 μm.

### Slide 22
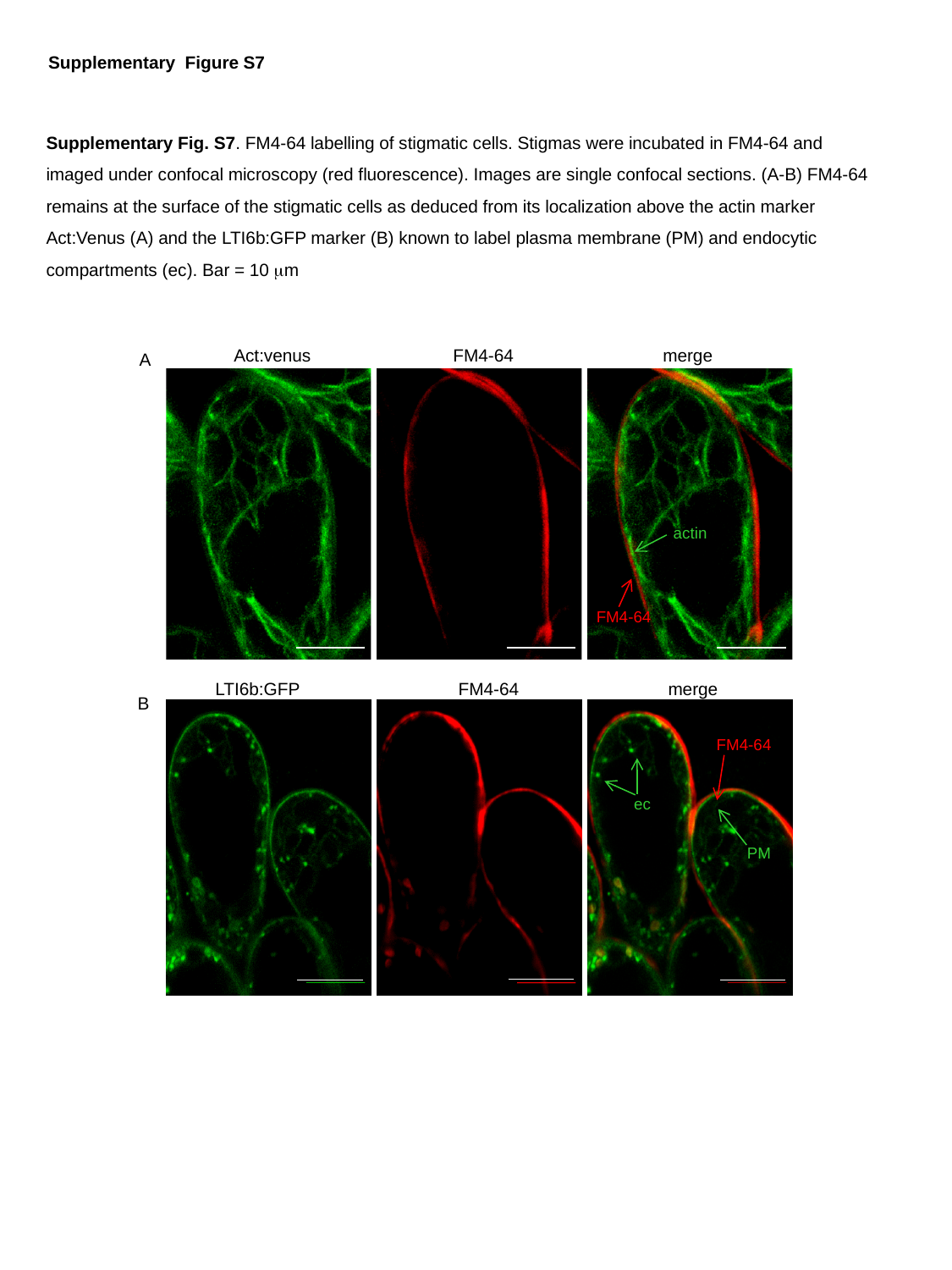

Supplementary Figure S7
Supplementary Fig. S7. FM4-64 labelling of stigmatic cells. Stigmas were incubated in FM4-64 and imaged under confocal microscopy (red fluorescence). Images are single confocal sections. (A-B) FM4-64 remains at the surface of the stigmatic cells as deduced from its localization above the actin marker Act:Venus (A) and the LTI6b:GFP marker (B) known to label plasma membrane (PM) and endocytic compartments (ec). Bar = 10 μm
Act:venus
FM4-64
merge
A
actin
FM4-64
LTI6b:GFP
FM4-64
merge
B
FM4-64
ec
PM
